# Structural atlas of *Pakpunavirus* P7-1 reveals determinants of virion stability and genome ejection

**DOI:** 10.1101/2025.11.26.690771

**Authors:** Fenglin Li, Nathan Bellis, Ravi K Lokareddy, Chun-Feng David Hou, Ruoyu Yang, Steven Branston, Zsuzsanna Kovach, Renae Geier, Angela Soriaga, Lucy Sim, Pierre Kyme, Deborah Birx, Sebastien Lemire, Gino Cingolani

## Abstract

Bacteriophages of the *Pakpunavirus* genus exhibit broad host range and potent bacteriolytic activity, making them promising candidates for clinical use. Here, we present a structural atlas of the therapeutic phage *Pakpunavirus* P7-1, a component of a phage cocktail targeting *Pseudomonas aeruginosa* that has undergone Phase 1/2 clinical trials. We determined the near-atomic structure of the extended virion and obtained a medium-resolution reconstruction of the contracted tail. Atomic models were built for 20 structural proteins comprising the icosahedral capsid, neck, contractile tail, and baseplate. We identified six upward-pointing Short Tail Fibers that stabilize the extended sheath and six highly flexible Long Tail Fibers likely involved in host recognition. Ordered fragments of the Tape Measure Protein revealed six copies inside the tail tube, forming a 3-helix cork at the tail tip. Sheath contraction repositions the baseplate, projecting all twelve tail fibers outward, yet contraction alone is insufficient to trigger genome ejection.

## INTRODUCTION

*Pseudomonas aeruginosa* is a major cause of illness and death worldwide. The bacterium creates biofilms that are resistant to many antibiotics and common antimicrobials ^1^. This problem is made worse by the rapid emergence and spread of *P. aeruginosa* strains that have acquired antibiotic-resistant genes. *P. aeruginosa* infections are common in cystic fibrosis (CF), a multi-organ disease ^2,3^ marked by repeated bacterial infections in the lungs and ongoing inflammation. Early in life, the microbial community mainly includes *Staphylococcus aureus*, and *P. aeruginosa* becomes prevalent in later stages ^4^. Strains from CF patients often mutate and develop adaptive responses to antibiotics, leading to multidrug-resistant strains ^5^. Phage therapy for *P. aeruginosa* is increasingly seen as a promising treatment for CF-related infections ^6,7^. Therefore, there is growing interest in understanding known lytic *P. aeruginosa* phages and discovering new ones as antimicrobials against both antibiotic-sensitive and multidrug-resistant strains ^8^. We previously shared a detailed structural analysis of *Pseudomonas phages* Pa193 ^9^ and Pa223 ^10^ from the Armata Pharmaceuticals collection.

This study describes the complete architecture of the *Pseudomonas* phage P7-1, a *Myoviridae* phage from the *Pakpunavirus* genus, characterized by a long contractile tail ^11^. P7-1 was originally isolated from sewer samples in Southern California by Armata Pharmaceuticals. It has a 93.7 kb long double-stranded DNA (dsDNA) genome that includes 198 predicted ORFs (Supplementary Table 1) and 16 tRNA genes. Because of its stability and lytic activity, P7-1 was chosen for inclusion in a candidate anti- *Pseudomonas* phage cocktail, AP-PA02, which has undergone phase 1/2 clinical trials for the treatment of patients with cystic fibrosis (SWARM-P.a., NCT04596319) or non-cystic fibrosis bronchiectasis (Tailwind NCT05616221) ^12^.

A growing number of new *Pakpunaviruses* have been isolated in recent years. The latest release of the International Committee on Taxonomy of Viruses (ICTV) ^13^ (release v3, June 19, 2024) includes 35 members of the *Pakpunavirus* genus. However, over 60 *Pakpunaviruses’* genomes are available in public databases and partially annotated. Typically, these phages have genomes ranging from 88 to 95 kb, about one-third larger than those of phages in the *Pbunavirus* genus, such as E217 ^14^ or Pa193 ^9^. The genome size of *Pakpunaviruses* is roughly half that of phage T4 ^15^, which has a genome of approximately 168.9 kbp, encodes about 300 gene products, and has been studied in much greater depth.

In this study, we use cryo-EM Single Particle Analysis (SPA), localized reconstruction ^16^, proteomics, comparative genomics, and AlphaFold-guided structure prediction ^17^ to generate a near-atomic atlas of P7-1 with the extended and contacted sheath. Our study identifies all the structural proteins of phage P7-1, including the multi-subunit baseplate complex. We uncover new features and conserved elements previously observed in other long-tailed bacteriophages, shedding light on the machinery and mechanisms of genome delivery in *P. aeruginosa* ^18,19^, and enhancing our understanding of *Pakpunavirus* biology.

## RESULTS

### Organization of *Pseudomonas* phage P7-1

We vitrified the P7-1 mature virion and collected a high-resolution dataset on a 300 kV Krios microscope (Table 1). Morphologically, the phage has a long contractile tail spanning more than 1,200 Å and a small baseplate (Fig. 1a, Supplementary Fig. 1a), sharing some similarities with *Pbunavirus* phages E217 ^14^ and Pa193 ^9^. Surprisingly, the vast majority of vitrified virions (>99%) had an extended tail and retained the genome within the capsid, in stark contrast to E217 ^14^, which has a contracted tail in about 30% of particles after vitrification. To visualize the contracted tail, we subjected P7-1 to prolonged incubation at pH 4.5 (Table 1), which triggered tail contraction in approximately half of the particles on the grid (Fig. 1b, Supplementary Fig. 1b). However, unexpectedly, only a few particles with a contracted tail also ejected the genome, suggesting that tail-contraction and genome ejection are not interdependent processes in this phage.

**Figure 1.**
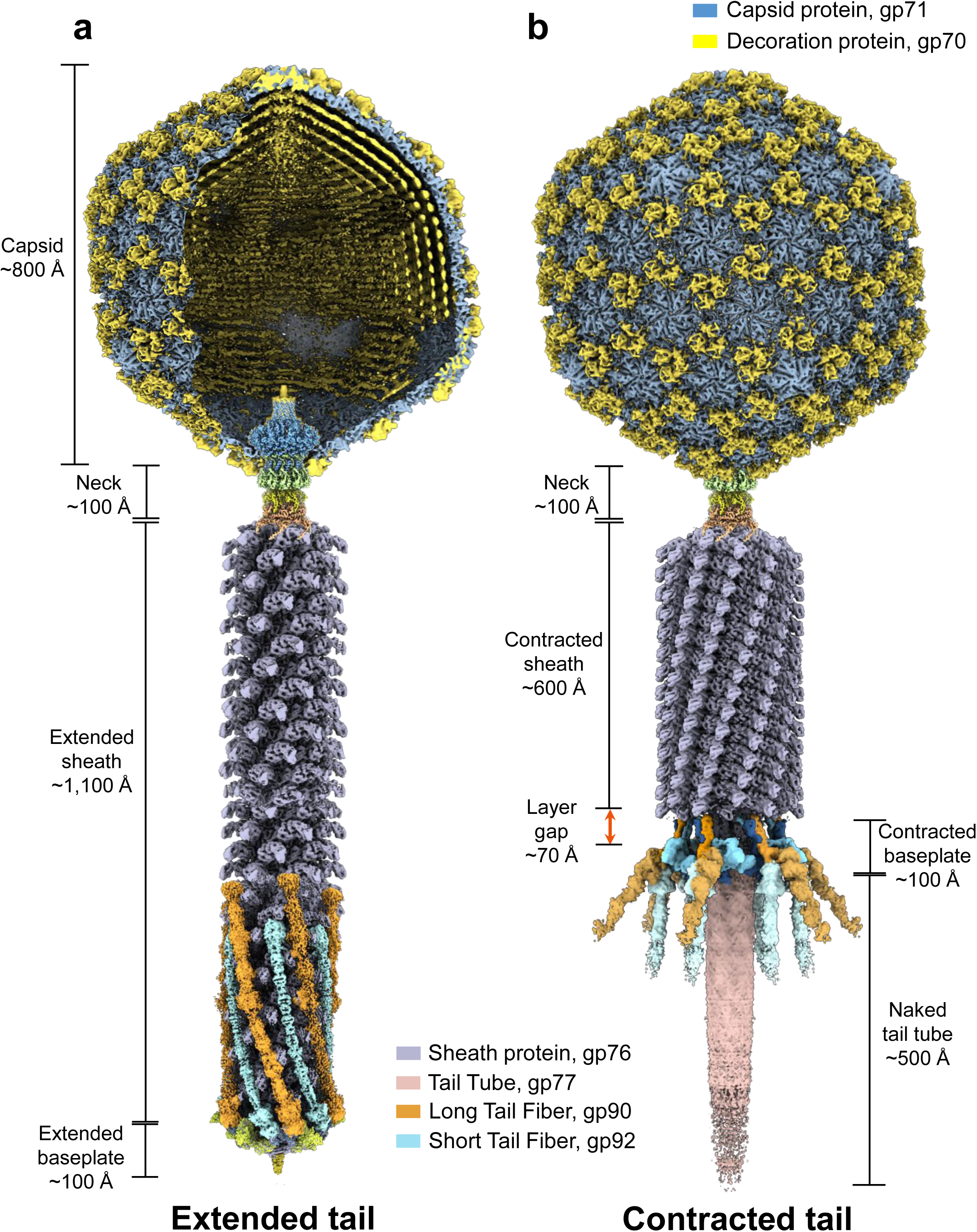
*Pseudomonas* phage P7-1 featuring an extended and contracted sheath. Composite cryo-EM reconstructions of (a) P7-1 virion at neutral pH with an extended sheath and (b) virion after incubation at acidic pH that triggers sheath contraction.

**Table 1.** Cryo-EM data collection and refinement statistics.

| Data Collection Statistics |  | P7-1<br>Extended<br>(pH 7.0) |  |  |  | P7-1<br>Contracted<br>(pH 4.5) |
| --- | --- | --- | --- | --- | --- | --- |
| Facility/Microscope |  | NCEF /<br>Titan Krios |  |  |  | S<sup>2</sup>C<sup>2</sup> /<br>Titan Krios |
| Camera |  | Gatan K3 |  |  |  | Falcon 4 |
| Magnification |  | X81,000 |  |  |  | X81,000 |
| Voltage (kV) |  | 300 |  |  |  | 300 |
| Electron exposure (e<sup>-</sup>/Å<sup>2</sup>) |  | 50 |  |  |  | 50 |
| Defocus range (um) |  | -1.2 to -2.4 |  |  |  | -1.2 to −2.4 |
| Pixel size (Å) |  | 0.681 (physical: 1.362) |  |  |  | 1.1 |
| Total movies (frames/movie) |  | 5,714 (40) |  |  |  | 10,329 (40) |
| Map and Model Refinement Statistics |  |  |  |  |  |  |
| PDB / Map entry codes |  | 9ZBI /<br>EMD-73989 | 9ZBS /<br>EMD-73999 | 9ZD8 /<br>EMD-74057 | 9ZD7 /<br>EMD-74056 | 9ZDK /<br>EMD-74065 |
| Particles per reconstruction |  | 12,148 | 12,148 | 10,400 | 10,400 | 7,832 |
| Map name |  | Capsid | Neck | Extended<br>Baseplate | Extended<br>Baseplate<br>with STF | Contracted<br>baseplate |
| Symmetry imposed |  | **C5** | **C6** | **C3** | **C3** | **C6** |
| Map resolution (Å) |  | 3.3 | 3.6 | 3.6 | 4.2 | 4.4 |
| FSC threshold |  | 0.143 | 0.143 | 0.143 | 0.143 | 0.143 |
| Models fit into density |  | Capsid,<br>decoration<br>proteins | Portal,<br>HT-adaptor,<br>collar,<br>gateway | Tube-B/-C,<br>ripcord, hub,<br>tip, adaptor-1/2, Triplex,<br>TMP | Tube-B/-C,<br>ripcord, hub,<br>tip, adaptor-1/2, Triplex,<br>TMP, STF | Contracted<br>Triplex,<br>adaptor-2,<br>sheath |
| Initial model (PDB code) |  | 8FRS | 8FVH | 8FUV, 8EON | 8FUV, 8EON<br>AlphaFold3 | 8FVG |
| Model-to-Map<br>Correlation Coefficient (mask) * |  | 0.82 | 0.85 | 0.79 | 0.83 | 0.80 |
| Model<br>composition | No. of chains | 9 | 36 | 63 | 81 | 30 |
|  | Non-H atoms | 19,323 | 72,366 | 107,973 | 174,997 | 80,631 |
|  | Residues | 2,409 | 9,006 | 13,938 | 22,895 | 10,527 |
| R.M.S.<br>deviations | Bond lengths (Å) | 0.017 | 0.003 | 0.003) | 0.003 | 0.002 |
|  | Bond angles (°) | 0.609 | 0.502 | 0.519 | 0.590 | 0.525 |
| Validation | MolProbity score | 1.78 | 1.72 | 2.40 | 2.47 | 2.61 |
|  | Clashscore | 4.05 | 5.25 | 6.55 | 8.42 | 11.61 |
|  | Rotamer outliers (%) | 5.30 | 1.50 | 7.24 | 8.26 | 11.14 |
| Ramachandran<br>plot | Favored (%) | 97.83 | 95.67 | 94.31 | 95.44 | 93.14 |
|  | Allowed (%) | 2.13 | 4.32 | 5.58 | 4.40 | 6.14 |
|  | Outliers (%) | 0.04 | 0.01 | 0.11 | 0.17 | 0.72 |

\* CC is calculated inside a mask around the model.

We used approximately 12,000 particles to reconstruct the entire mature virion of P7-1 with an extended tail, utilizing focused maps of the capsid, neck/tail, and extended baseplate (Table 1, Supplementary Figs. 2a-c). At the same time, about 7,800 tail-contracted, head-filled virions were used to determine a medium-resolution (∼4.2 Å) C6 reconstruction of the contracted tail (Table 1, Supplementary Fig. 2d), which revealed significant restructuring of the baseplate complex. Using the four maps described above, we built atomic models for all structural proteins in P7-1. The capsid was visualized using a C5 localized reconstruction at a resolution approaching 3.3 Å (Supplementary Fig. 2a, e). The neck and tail were reconstructed using localized reconstruction by selecting a class of particles with tails at one five-fold vertex and imposing C6 symmetry, which yielded a 3.6 Å map (Supplementary Fig. 2b, e). Finally, we reconstructed the baseplate from the extended tail by manually picking 10,400 baseplate particles and applying C6 and C3 symmetry to obtain reconstructions at 3.6 Å (Supplementary Fig. 2c, e). The contracted baseplate was modeled in a C6 4.2 Å density (Supplementary Fig. 2d, e).

All reconstructions exhibited strong density features, which we utilized to annotate 19 P7-1 Open Reading Frames (ORFs) and generate atomic models for 20 full-length gene products (gp), as illustrated in Fig. 2a-c. The structural atlas of P7-1 presented in this paper includes three capsid proteins (gp67, gp70, and gp71) (Fig. 2a), three neck proteins (gp72, gp74, and gp75), two tail proteins (gp76 and gp77) (Fig. 2b), and eleven baseplate proteins (gp78, gp79, gp83, gp84, gp85, gp86, gp87, gp88, gp89, gp90, and gp92) (Fig. 2c). The large Triplex component gp88 exists in two conformations (gp88-a and gp88-b). All atomic models built into cryo-EM maps were real-space-refined to achieve a model-to-map correlation coefficient (CC) between 0.79 and 0.85 (Table 1), except for the long tail fibers, which have weak density and were not refined (described below). Finally, we also modelled two small regions of the Tape Measure Protein (TMP) that are not shown in Figure 2. All structural gene products annotated using cryo-EM were also identified and validated by liquid chromatography-mass spectrometry (LC-MS) of P7-1 virions (Supplementary Table 2), with one exception: gp68/gp69, which are likely located inside the capsid and are invisible by SPA.

**Figure 2.**
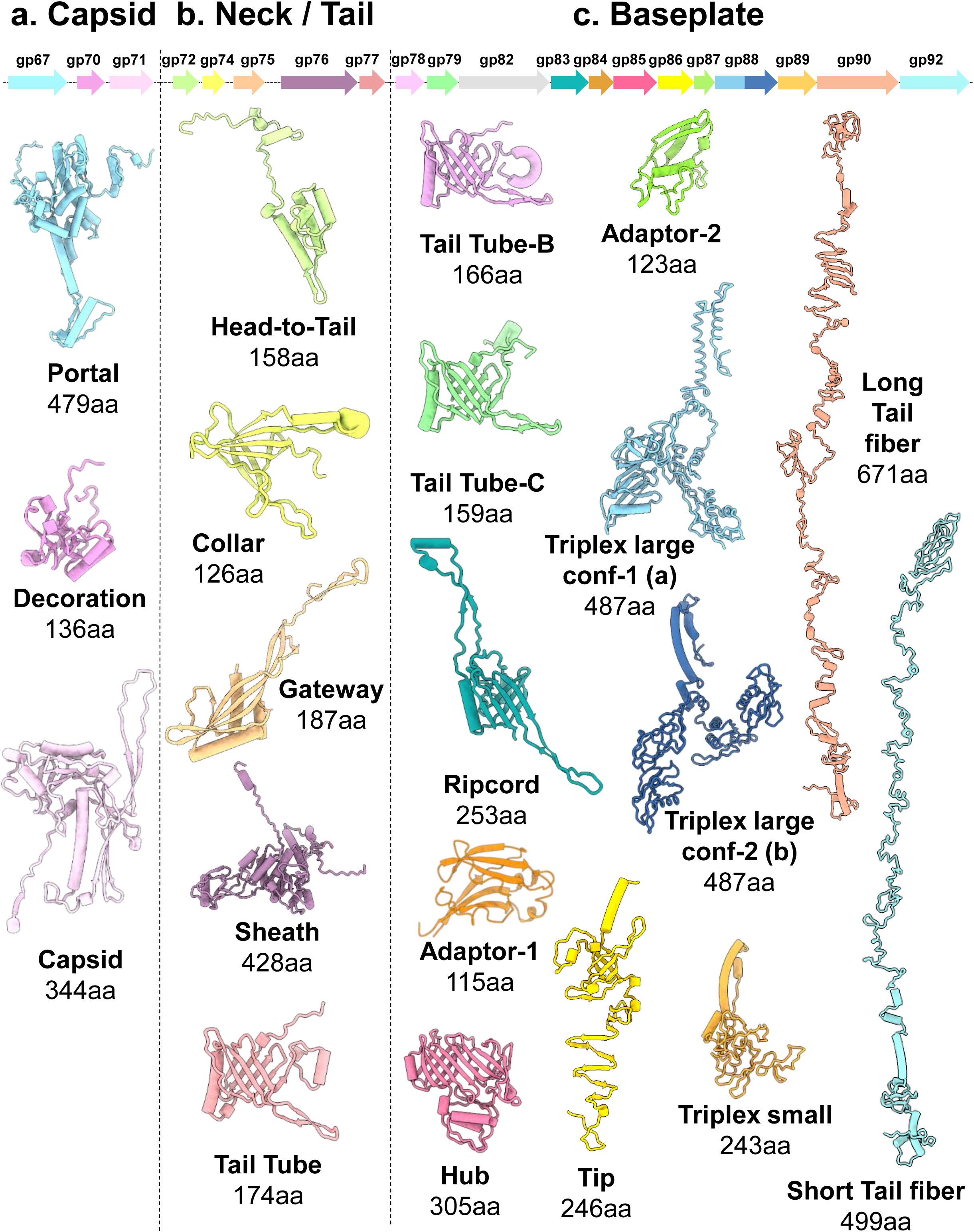
Structural atlas of phage P7-1 structural proteins. Ribbon diagrams of 20 gene products were identified and built *de novo* in this study. The gene products are divided into three groups: (a) three capsid factors; (b) three neck factors and two tail factors; (c) eleven gene products that are part of the baseplate, encoded by ten genes. The Triplex large subunit factor gp88 exists in two conformations: conformation-1 (gp88-a) and conformation-2 (gp88-b). Fragments of the TMP (gp82) are not displayed in this figure.

### The capsid and neck of *Pseudomonas* phage P7-1

P7-1 capsid is large, with a maximum diameter of approximately 800 Å. A 3.3 Å focused reconstruction of the P7-1 capsid’s five-fold vertex (Supplementary Fig. 2a) revealed two structural components: the major capsid protein gp71, which forms an icosahedron with a triangulation number T = 9, and the decoration protein gp70 (Fig. 1a, 2a, Table 1). Gp71 (344 aa) has a canonical HK97 fold, most similar to the major capsid protein of the flagellotropic phage YSD1 ^20^, a *Siphoviridae* (E-value = 2.32e-15; RMSD = 5.52 Å, 344 versus 354 residues in coverage). Gp70 folds into a homotrimer (Fig. 1a, 2a) that is topologically superimposable to phage YSD1 cementing protein ^20^ and overall very similar to the prototypical head decoration protein gpD from phage lambda ^21^. One hundred eighty copies of the trimeric gp70 bind the capsid exterior at each local three-fold axis (Fig. 1a,b), generating a second capsid layer that decorates ∼60% of the exterior surface of the P7-1 capsid. A similar pattern was observed in the *Pseudomonas* phages E217 ^14^ and Pa193 ^9^, consistent with gp70 functioning as a putative cementing protein.

We generated C12 and a C6 symmetric reconstructions of phage P7-1 unique vertex that revealed the architecture of the phage neck and the beginning of the extended tail (Supplementary Fig. 2b). The C12 map was used to identify and build *de novo* the portal protein gp67 and the head-to-tail (HT) adaptor gp72 (Fig. 2b, 3a, Table 1). These two proteins form a dodecameric complex at one of the 12 five-fold vertices of the capsid, replacing a penton. The portal interacts with the capsid protein (gp71) via a 12:10 symmetry-mismatched binding interface ^22^. The P7-1 portal protein is relatively small, consisting of only 479 amino acids, compared to 722 residues in the podophage P22, for example ^23,24^. We built a nearly complete structure (res. 10-454) in a 3.6 Å map, while no density was visible for the first nine and last 25 amino acids. The P7-1 portal protein is most similar to the portal protein from bacteriophage G20C ^25^ (E-value = 1.67e-17), with 20.1% sequence identity and a comparable length (427 versus 479 residues). It features a classical portal protein fold consisting of crown, wing, and stem domains, but notably lacks a C-terminal barrel (Fig. 3a, b) ^22^, possibly explaining its smaller size. P7-1 HT-adaptor ring assembles co-axially under the portal, wrapping around the portal clip and stem regions ^23^. The HT-adaptor consists of an N-terminal 4-helix bundle core with an extended C-terminal arm that contains three α-helices wrapped around the portal clip and stem regions, resembling a bouquet wrap (Fig. 3a,b). HT-adaptor binding to the portal protein during genome packaging competes with the large terminase subunit (TerL), ending the packaging reaction ^26^.

**Figure 3.**
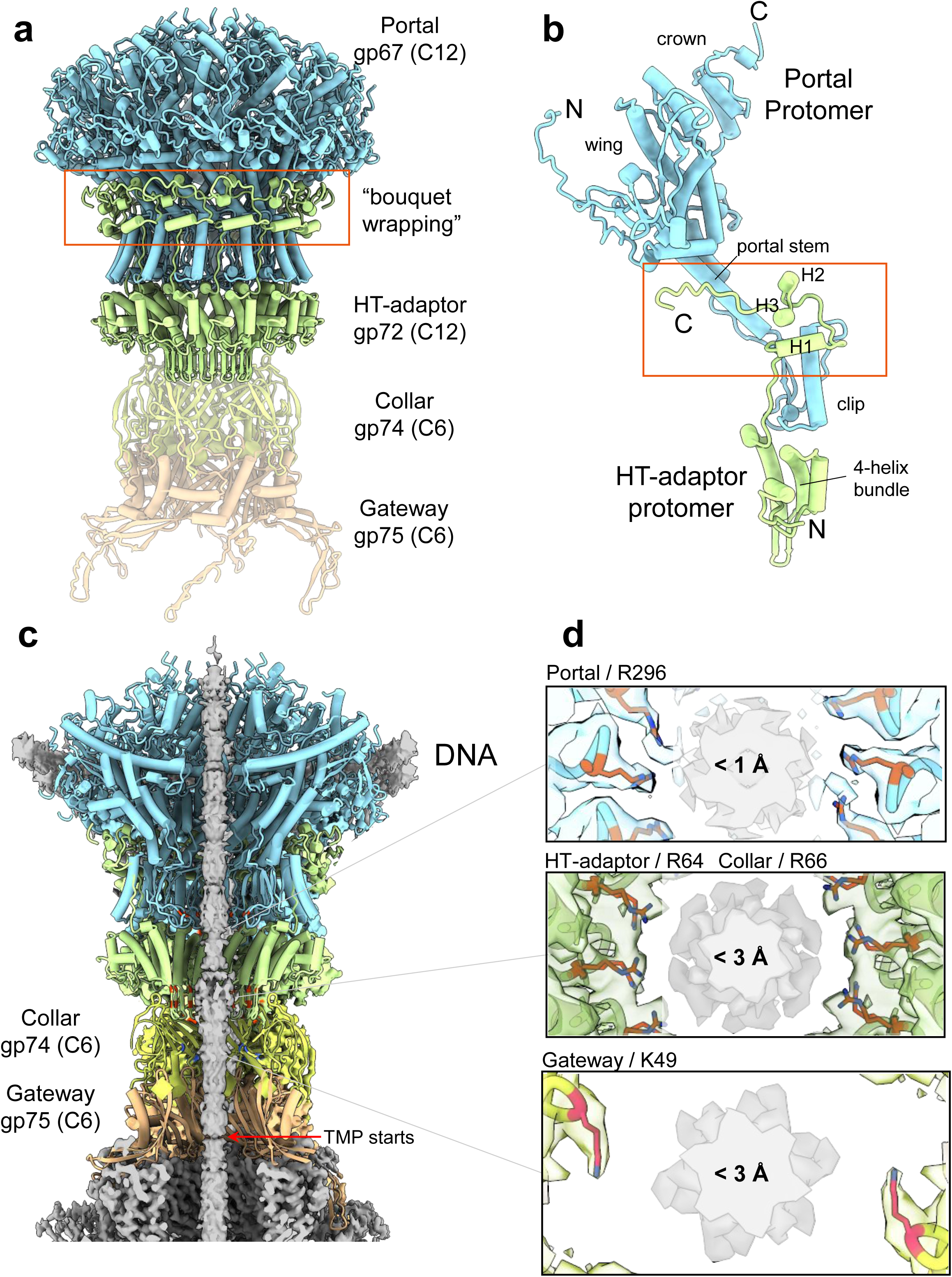
Structure of phage P7-1 neck assembly. (a) Overview of the P7-1 neck complex. The dodecameric portal protein gp67 (cyan) binds to twelve copies of the HT-adaptor gp72 (light green), which then binds to the collar gp74 (light yellow) and the gateway gp75 (light orange). (b) A ribbon diagram of a portal protein monomer bound to the HT-adaptor reveals the bouquet-wrapping interaction (red frame) between the C-terminal moiety of the HT-adaptor and the portal clip and stem regions. (c) A cut-out view of the P7-1 neck, as shown in panel (a), reveals a density that extends along the entire length of the channel. (d) The top view of the neck channel illustrates the interaction between the neck component and the density filling the channel (likely dsDNA) across three distinct regions: portal, HT-adaptor/collar, and gateway.

All P7-1 factors under the HT-adaptor are hexameric ^27^ and were built in a C6 symmetry map (Supplementary Fig. 2b), specifically, the collar gp74 and gateway gp75 that extend the portal:HT-adaptor channel by approximately 80 Å (Fig. 3c). Notably, the neck and tail channel is filled with a density, possibly double-stranded DNA, which is C6 averaged in our reconstruction. Additionally, the internal diameter of the P7-1 channel varies from portal to gateway. The portal channel and HT-adaptor inner cavities are narrow, with close contacts (< 4 Å) existing between basic residues protruding into the channel (e.g., portal R296; HT-adaptor R64, etc.) and the inner density, suggesting that the internal density corresponds to DNA (Fig. 3d). In contrast, the gateway cavity at its widest point is significantly larger, approximately 45 Å between two juxtaposed residues K49, suggesting energetically weaker contacts between the tail tunnel and the density inside the channel. The density in this part of the inner channel appears bulkier (Fig. 3d) and does not match dsDNA, suggesting it is probably the TMP, as identified by LC-MS analysis (Supplementary Table 2).

### P7-1 tail and baseplate proteins

The P7-1 tail consists of a repetitive set of tail tube hexamers surrounded by sheath protein subunits arranged in a helical pattern. We constructed P7-1 with an extended tail approximately 1,100 Å long, comprising 204 copies of tail tube gp77 (residues 1-174) and 204 copies of tail sheath gp76 (residues 1-428) ^28^ (Figs. 1a, 2a). The tail tube has 22.5% sequence identity to the Cyanophage Pam3 tube ^29^ (with comparable amino acid lengths of 174 versus 142 residues) and a similar 3D-fold (RMSD of 2.1 Å). The extended tail consists of an internal component comprising 34 stacks of hexameric tube protein with a lumen approximately 37 Å wide. P7-1 sheath protein gp76 (residues 1-428) has a three-domain architecture similar to E217 ^14^, but it differs from the Pam3 sheath protein, which contains only two domains (Supplementary Figure 3a,b). N- and C-terminal extensions connect sheath proteins to neighboring subunits, allowing the protein to assemble around the tube structure in a helical arrangement. The topology of the sheath assembly and the conformational changes that occur upon contraction are similar to those described for other contractile *Myoviridae* ^9,14^ and will not be discussed here.

The separate reconstruction of the P7-1 extended baseplate, determined at resolutions of 3.6-4.5 Å (Table 1, Supplementary Fig. 2c,e), allowed us to identify and build atomic models for 11 polypeptide chains (Fig. 2c), each present in 63 copies. The baseplate complex can be divided into three subcomponents, organized into a metastable assembly poised for contraction ^30^. First, the baseplate cap subcomplex (Fig. 4a), which remains stationary during sheath contraction, is made up of a highly complex extension of the repetitive tail tube. The P7-1 cap subcomplex consists of two tail tube-like subunits (tail-tube-B gp78 and tail-tube-C gp79, which are 25.5% identical), linked by the ripcord (gp83) and hub (gp85) subunits to a trimeric tail tip (gp86) (Fig. 2c), which seals the tail channel. The tail tube variants gp78 and 79, and the ripcord form hexameric rings that are coaxial to the tail tube gp77. In contrast, the hub and tail tip are trimeric (Fig. 4a). The P7-1 tail tip features a classical triple β-helix fold ^31^, which is also found in analogous contractile ejection systems like Pa193 ^9^, Pam3 ^29^, R-type pyocin ^32^, and E217 ^14^. The second component of the P7-1 baseplate consists of two adaptor subunits, gp84 (adaptor-1, 115 residues) and gp87 (adaptor-2, 123 residues) (Fig. 2c), which bind to the outward surface of the tail hub (gp85) and ripcord (gp83), respectively (Fig. 4b). The adaptor subunits decorate the cap complex without making direct contact with one another, and thus does not form an oligomer like all the other factors of the cap complex. Finally, the P7-1 baseplate is built by a nut-subcomplex (Fig. 4c) that slightly expands to a maximum diameter of ∼290 Å at the tail tip distal to the capsid. The nut-subcomplex comprises six copies of the Triplex complex (Fig. 4d), attached like a nut to a bolt. Each Triplex complex consists of two copies of the large subunit gp88 in structurally distinct conformations, gp88-a and gp88-b, along with one copy of the small subunit gp89 (Fig. 2c). The three subunits combine to form an extended helical bundle that contributes to much of the association energy holding the complex together. The nut-subcomplex is likely metastable in the extended tail conformation, ready for upward motion during sheath contraction ^14^.

**Figure 4.**
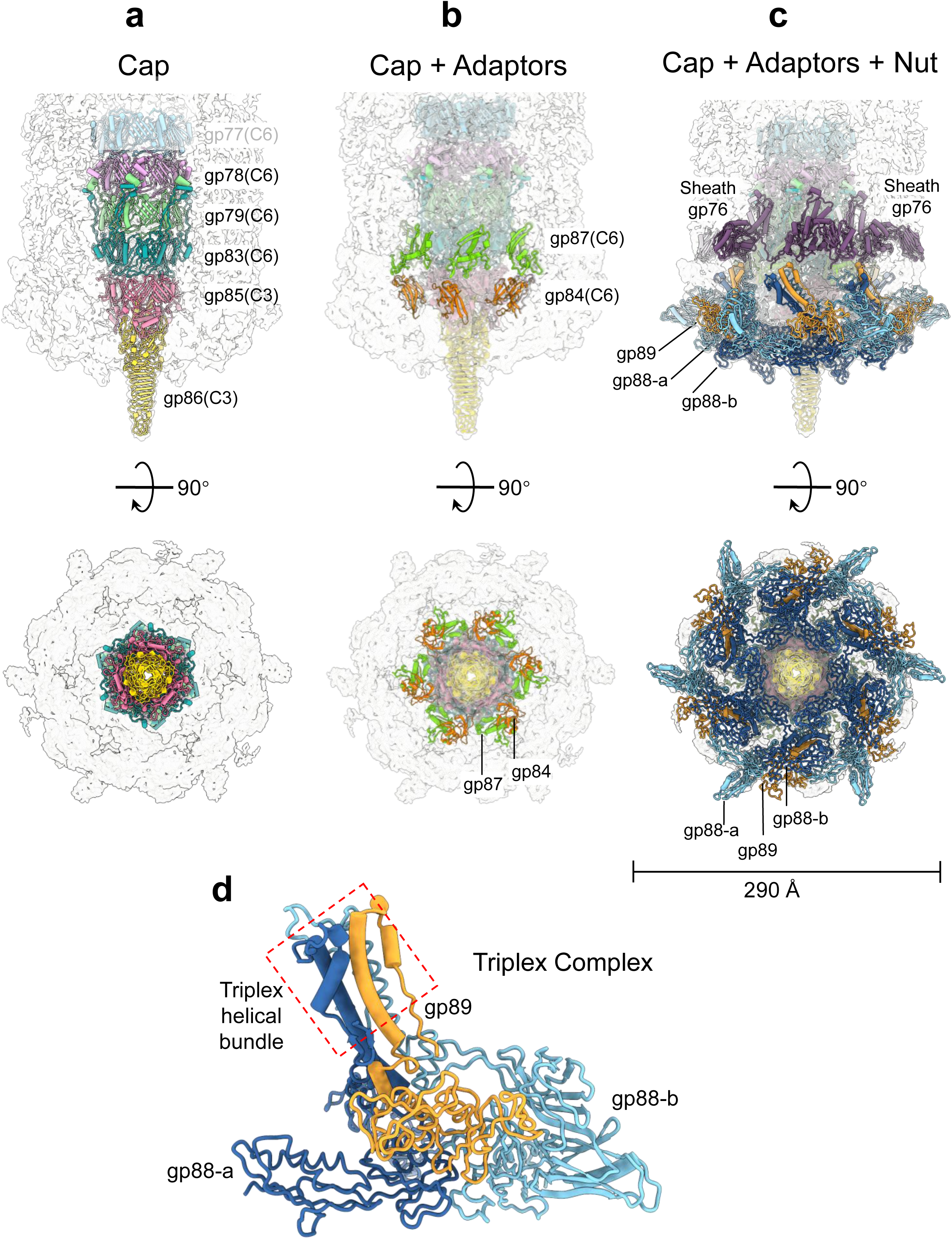
Anatomy of phage P7-1 baseplate. (a) A semitransparent map of the P7-1 baseplate, overlaid with ribbon diagrams of the baseplate cap, which includes one oligomer of the tail tube gp77 and three coaxial tube-like factors: gp78 (tail tube-B), gp79 (tail tube-C), gp83 (ripcord), tail hub gp85, and tail tip gp86. (b) The baseplate adaptor proteins, gp84 (adaptor-1) and gp87 (adaptor-2). (c) The baseplate nut complex consists of six copies of the triplex complexes (gp88-a, gp88-b, and gp89). For panels (a-c), the maps are shown in a side view and rotated by 90 degrees. (d) Ribbon diagram of P7-1 triplex complex formed by gp88-a, gp88-b, and gp89.

### P7-1 contains two Tail Fiber proteins

The C3 reconstruction of the P7-1 baseplate revealed prominent density features that point upward, visible at a high contour, spiraling approximately 400 Å along the tail and contacting the sheath lattice (Fig. 5a, left). We assigned this density to the Short Tail Fiber (STF) gp92 (Fig. 2c), which we modeled using AlphaFold3 ^33^ as a slender trimer of 499-residue subunits. We docked the STF model into the density and real-space-refined it against a focused reconstruction of the baseplate to CC = 0.83 at 4.2 Å resolution (Table 1). STF measures approximately 410 Å in length and about 25 Å in thickness over most of its span (Fig. 5b). It features an N-terminal globular knob that is roughly 40 Å in diameter, as well as a C-terminal knob that is approximately 60 Å long (Fig. 5c). By lowering the map contour (Fig. 5a, right), we also noticed a second elongated density pointing upward along the tail sheath lattice, roughly parallel to STF. We attributed this density to the Long Tail Fiber (LTF) gp90, which comprises 671 amino acids (Fig. 2c). Although the LTF density is much weaker than that of STF, an AlphaFold3 model of trimeric gp90 fits surprisingly well within the C3 cryo-EM map (Fig. 5a, right). Six copies of LTF were docked per baseplate, which, together with six STF, yield twelve tail fibers (Fig. 1a). Unlike STF, which refined well against the C3 density (CC = 0.83 at 4.2 Å, Table 1), the LTF model was not refined against the weak density to prevent overfitting. Comparing the two fibers reveals interesting differences. Both fibers are slightly bent, but LTF is longer (480 Å in length) and spirals as it meets the tail attachment site at the tail tip (Fig. 5c). It contains five knobs: three internal (I-knob-1, I-knob-2, and I-knob-3) and two at the opposite N- and C-terminal tips, each approximately 40 Å in diameter (Fig. 5c). Both STF and LTF attach to the Triplex complex small subunit, gp89 (Fig. 5d). This subunit specifically contains two fiber attachment loops (FAL), FAL1 (res. 92-112) and FAL2 (res. 215-236) (Fig. 5e), which act as flexible anchoring surfaces for STF and LTF, respectively.

**Figure 5.**
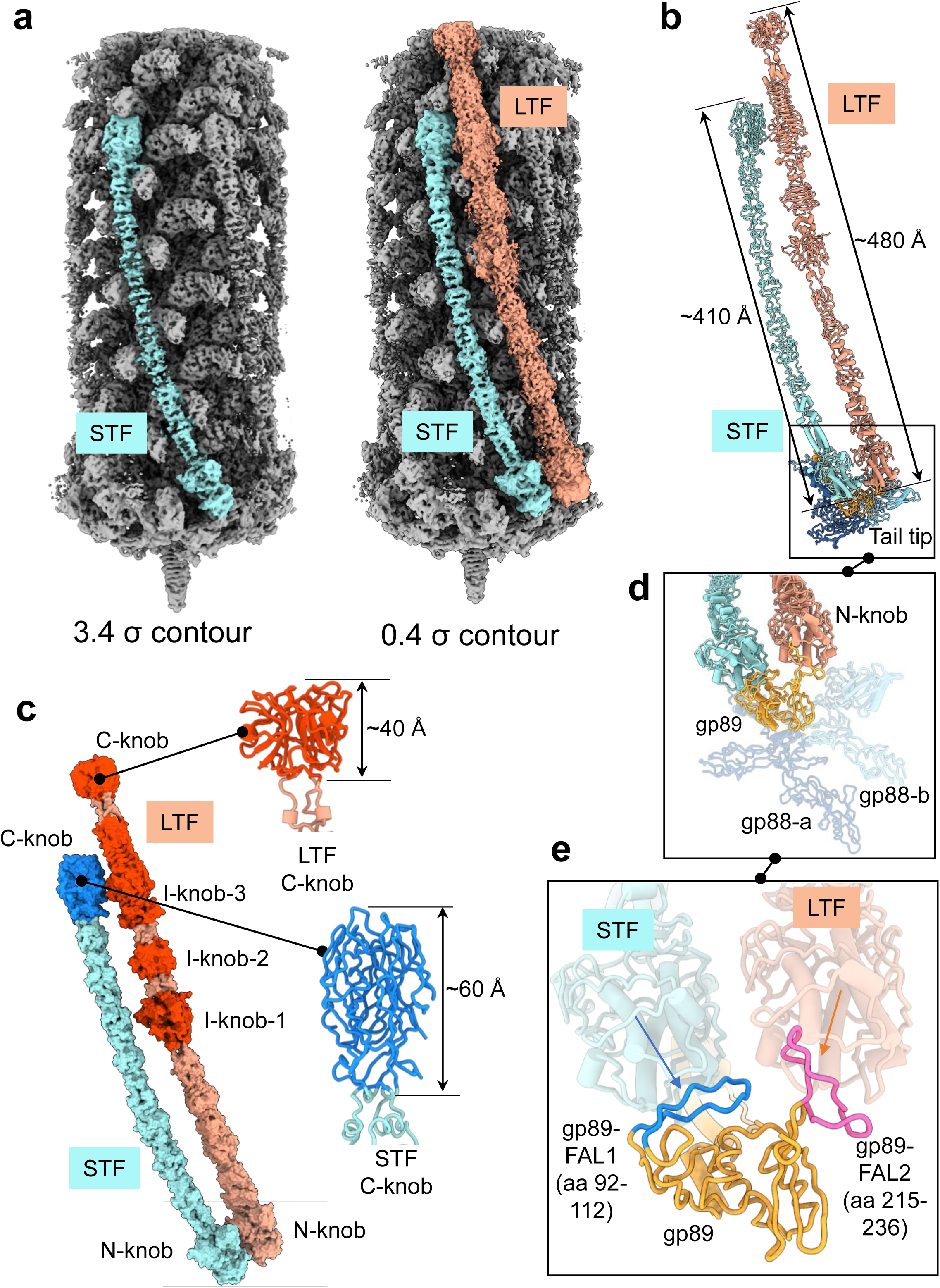
Structures of P7-1 tail fibers and their attachment to the baseplate. (a) A semi-transparent C3 localized map of the P7-1 baseplate and tail, computed at 4.2 Å resolution, is contoured at 3.4σ (left) and 0.4σ (right). STF (cyan) is visible at both contours, while density features for LTF (salmon) are only detectable at the lower contour. (b) Cartoon schematic of the isolated STF and LTF assembled with a Triplex complex. (c) The solvent-surface representation of STF and LFT highlights the positions of the knobs. STF contains two knobs, while LTF has five knob-like motifs: two at the N- and C-terminal tips and three internal (annotated as I-knob-1/2/3). Magnified views of STF and LTF C-knobs reveal the differing bulkiness of these two motifs. (d) Magnified view of the interface between the baseplate Triplex complex (gp89, gp88-a, gp88-b) and the STF/LTF N-knobs. (e) A zoomed-in view of gp89 fiber attachment loops FAL1 (blue) and FAL2 (magenta) that bind to STF and LTF n-knobs, respectively.

The stronger electron density seen in the cryo-EM reconstruction for STF compared to LTF (Fig. 5a) suggests that STF primarily remains in a closed conformation, stabilizing the extended tail. In contrast, LTF is looser and likely accounts for the fibers extending outward from the baseplate in the cryo-micrographs (Supplementary Fig. 1a). We attribute the enhanced rigidity of STF to three factors. <u>First</u>, the STF N-termini obtain additional supporting stability from an extensive binding interface with the Triplex helical bundle (Fig. 6a), which is not observed for the LTF (Fig. 6b). <u>Second</u>, only the STF inserts into the sheath lattice, which likely explains the higher sequence conservation of STF versus LTF, as it must make extensive contacts throughout its 210 Å length (Fig. 6c). Each sheath subunit makes two salt bridges, nine hydrogen bonds, and 198 nonbonded contacts with the STF, whose slender surface facilitates bonding with the sheath lattice. These interactions may not be as strong for the LTF, where three bulky I-knobs (Fig. 5c) sterically hinder bonding with sheath subunits. <u>Finally</u>, the STF C-knob is longer and bulkier than its counterpart in the LTF (Fig. 5c), allowing it to fit snugly within five layers of sheath molecules and anchor itself against the fifth layer (Fig. 6c).

**Figure 6.**
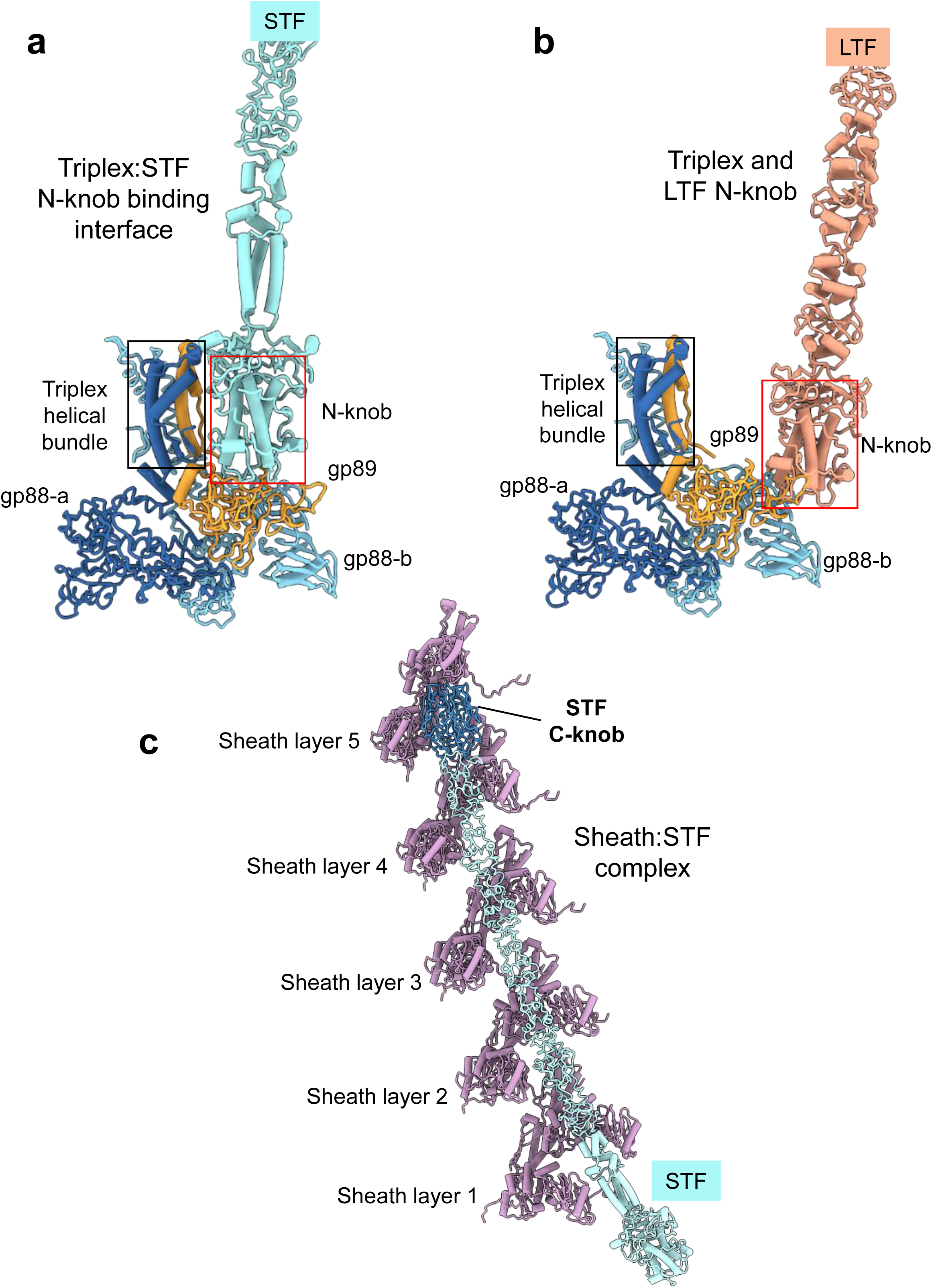
Determinants of P7-1 STF rigidity. (a) A ribbon diagram of the Triplex complex is shown next to the STF (a) and LTF (b). Only the STF N-knob makes close contact with the triple helical bundle (panel a), while the LTF N-knob is too distant to bond (panel b). (c) Ribbon diagram of STF and the five layers of sheath proteins. STF interacts with sheath subunits from 5 different layers within the P7-1 tail. The STF C-knob is colored a darker blue and stacks extensively against the 5^th^ sheath layer.

### Evolution of P7-1 tail fibers

Phage P7-1 carries 12 tail fibers arranged around the tail tip, distal to the capsid (Fig. 1a). Although they appear similar, the two fiber types are evolutionarily, structurally, and likely functionally distinct. Comparative analysis of structural protein-coding sequences from 61 *Pakpunavirus* genomes (Supplementary Table 3) revealed that STFs are highly conserved, with 92.4–99.6% identity across the *Pakpunavirus* genus (Supplementary Fig. 4). In contrast, LTFs are more variable, with ∼85% sequence identity overall (Supplementary Fig. 4). Domain-level analysis (Fig. 7a) revealed that the N-terminal region of LTF (residues 1–324), which attaches to the baseplate, is the most conserved (86.4–100%), while the middle and C-terminal regions are considerably more divergent.

**Figure 7.**
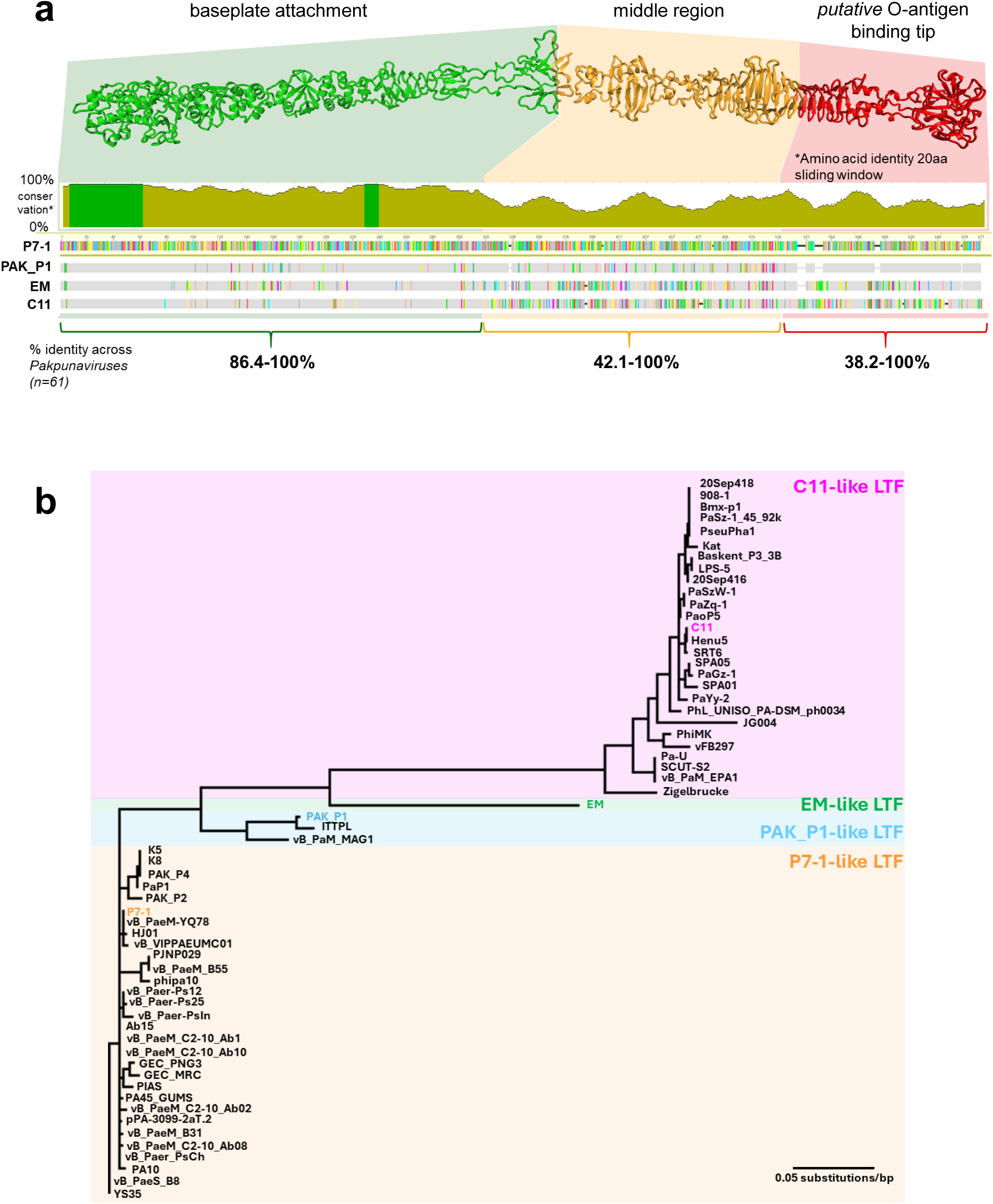
Evolutionary analysis of LTF in *Pakpunavirus*es. (a) A diagram showing 7-1 LTF (gp90) conservation (bottom) mapped onto its structure (top), divided into three regions. Each color represents an amino acid, with identical amino acids in phage 7-1 LTF shown in gray. Colored bars highlight discrepancies with the 7-1 LTF sequence. The N-terminal region (amino acids 1-324; green) is highly conserved across all phages. The middle segment (amino acids 325-540; orange) differs from the other three protein groups in phage 7-1. The C-terminal end (amino acids 541-671; red) is very similar between 7-1-like and PAK_P1-like phages but differs significantly between these groups and the EM-like and C11-like phages (Supplementary Table 4). (b) Phylogenetic tree of LTFs from 60 P7-1-like phages, showing four distinct clusters represented by P7-1 (yellow), PAK_P1 (cyan), EM (gray), and C11 (pink).

To evaluate whether LTF conservation reflects phage evolutionary subgroups, we performed a phylogenetic analysis of 60 publicly available *Pakpunavirus* genomes, excluding phage LCK69 due to an intron-split LTF gene (Supplementary Table 3). This analysis identified four phylogenetic groups (Fig. 7b; Supplementary Table 4). Twenty-nine phages, including P7-1, cluster together with >95.4% sequence identity. A second group of 27 phages, represented by C11, shares 91% identity among themselves but only 64.7–69.1% with other groups. C11-like and P7-1-like LTFs are highly conserved across residues 1–324 but diverge substantially thereafter (Fig. 7a; green vs. orange/red regions). A third group of three phages, represented by PAK_P1, shows >93.3% identity among themselves, ∼88% identity with P7-1-like LTFs, ∼67% identity with C11-like LTFs, and 74% identity with EM. PAK_P1-like LTFs retain strong conservation in the N-terminal domain (res. 1–324; Fig. 7a, green) and in the C-terminal domain (>93% identity to P7-1-like LTFs; Fig. 7a, red), but differ markedly in the intervening region (res. 325–540; Fig. 7a, orange). Finally, phage EM represents a fourth, more divergent lineage, with its LTF distinct from all others except for limited conservation at the N-terminus (residues 1–80).

Consistent with these sequence-based patterns, preliminary data from Armata Pharmaceuticals’ proprietary phage collection suggest that LTF C-terminal domains contribute to host specificity. In particular, a P7-1-like C-terminus is associated with infection of *P. aeruginosa* strains expressing O3, O4, or O11 O-antigens, while a C11-like C-terminus correlates with infection of O2-expressing strains (*unpublished data*). These findings support a central role for the LTF C-terminal region in determining host range in *Pakpunaviruses*. The contribution, if any, of the variable middle region (orange in Fig. 7a) remains unclear.

### Sheath contraction restructures the P7-1 baseplate

Prolonged incubation of P7-1 at acidic pH causes irreversible sheath contraction in approximately half of the phages on a cryo-grid (Supplementary Fig. 1b), which we used to determine a medium-resolution reconstruction of the contracted baseplate (Fig. 1b, Table 1). Even with its modest resolution, this C6 density proved helpful in modeling the contracted conformation of the P7-1 baseplate (Fig. 8a). Sheath contraction leads to the dissociation of the baseplate into two components, as seen in Twort-like *Myoviridae* phages ^34^. The baseplate cap subcomplex (comprising subunits gp78, gp79, gp83, gp85, and gp86) remains at the distal tail tip. The resolution of our reconstruction is too low to model these subunits. However, the fact that most contracted phages retain dsDNA in their heads (Supplementary Fig. 1b) strongly suggests that the tail tip continues to seal the tail after sheath contraction, and tail sheath contraction alone is not sufficient to induce genome release. In contrast, both the baseplate nut subcomplex (six copies of subunits gp88, gp89-a, and gp89-b) and six copies of the adaptor-2 (gp87) move upwards along the tail tube like a bolt around a nut (Fig. 8a). The adaptor-2 subunits, which decorate the outer surface of the ripcord complex in the extended baseplate (Fig. 4b), move upward to insert at the interface between sheath subunits, from the first layer of the contracted sheath. We modeled the N- and C-terminal moieties of the sheath from neighboring subunits that create a 4-stranded β-sheet with a β-hairpin from the adaptor-2 subunit, which effectively functions as a sheath terminator (Fig. 8c). A distinctive ∼70 Å gap is visible between the first sheath layer and the contracted baseplate (Fig. 1b, 8a), corresponding to the height of the Triplex complex helical bundle that bonds adaptor-2 subunits (Fig. 8b). Presumably, the binding of the tail fibers, especially LTF, to their receptors pulls the FALs downward, enabling a conformational change in the triplex that releases it from the tail tube termination subcomplex and allows the sheath to contract, bringing the nut-bolt subcomplex with it. (Fig. 8d). FAL1, the attachment site for STF, swings by 270°, while the more flexible LTF connecting loop, FAL2, swings by approximately 90°, projecting both short and long TFs downwards in the contracted state (Supplementary Fig. 5). Due to their flexibility and the lack of physical restraints in the contracted state, only about 200 amino acids of each fiber are visible in the reconstruction.

**Figure 8.**
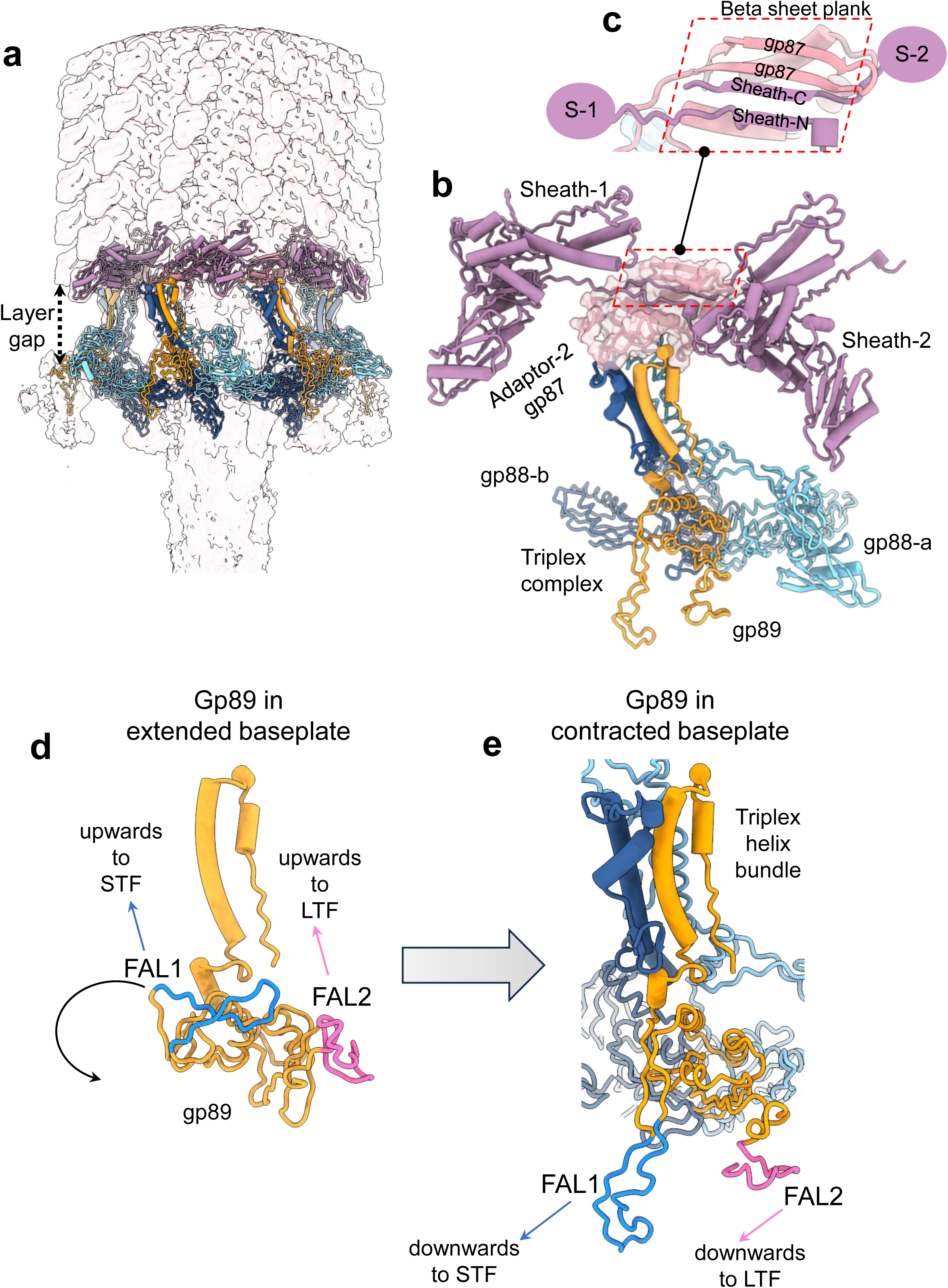
Conformational changes during sheath contraction. (a) The density map of the contracted baseplate, calculated at 4.2 Å, is displayed as a semi-transparent surface at a 2.5 σ contour level. Only six sheath proteins (purple) from the first layer and six copies of the Triplex complex are overlaid onto the density. (b) A magnified view reveals one Triplex complex, which is bonded to the adaptor-2 (gp87) (light pink), positioned between two sheath subunits. (c) The adaptor-2 forms a 4-stranded β-sheet with N- and C-terminal ends from neighboring sheath subunits. (d-e) Ribbon diagram of the Triplex small subunit gp89 before and after contraction. FAL1 and FAL2 are colored blue and magenta, respectively, and they swing downwards after contraction.

### Ordered regions of the TMP are visible within the cap subcomplex

The P7-1 tail tube is filled with density, likely consisting of the Tape Measure Protein (gp82), which is abundant in the MS analysis (Supplementary Table 2). By using a C3 map of the extended tail, we identified four stretches of ordered electron density spanning nearly 1,000 Å of the tail lumen, named as (i)-(iv) in Figure 9a. The first and most ordered region of density is located within the nut-subcomplex lumen at the interface with the tail tip as well as slightly above it, although lacking continuity (Fig. 9a(i), b). The density was interpreted using ModelAngelo ^35^ and found to correspond to the C-terminus of P7-1 TMP, residues 754-784. As recently observed in phage Pa193 ^9^ and previously reported for *Siphoviridae* phage HK97 ^36^, TMP C-terminal residues form a trimeric coiled-coil that sits like a cork onto the C-terminus of the tail tip, which is also arranged into a trimeric coiled-coil (Fig. 9c). We will refer to this part of TMP as the 3-helix cork. The cork is closely connected to the tail tip, which explains why the density is so well-ordered and traceable.

**Figure 9.**
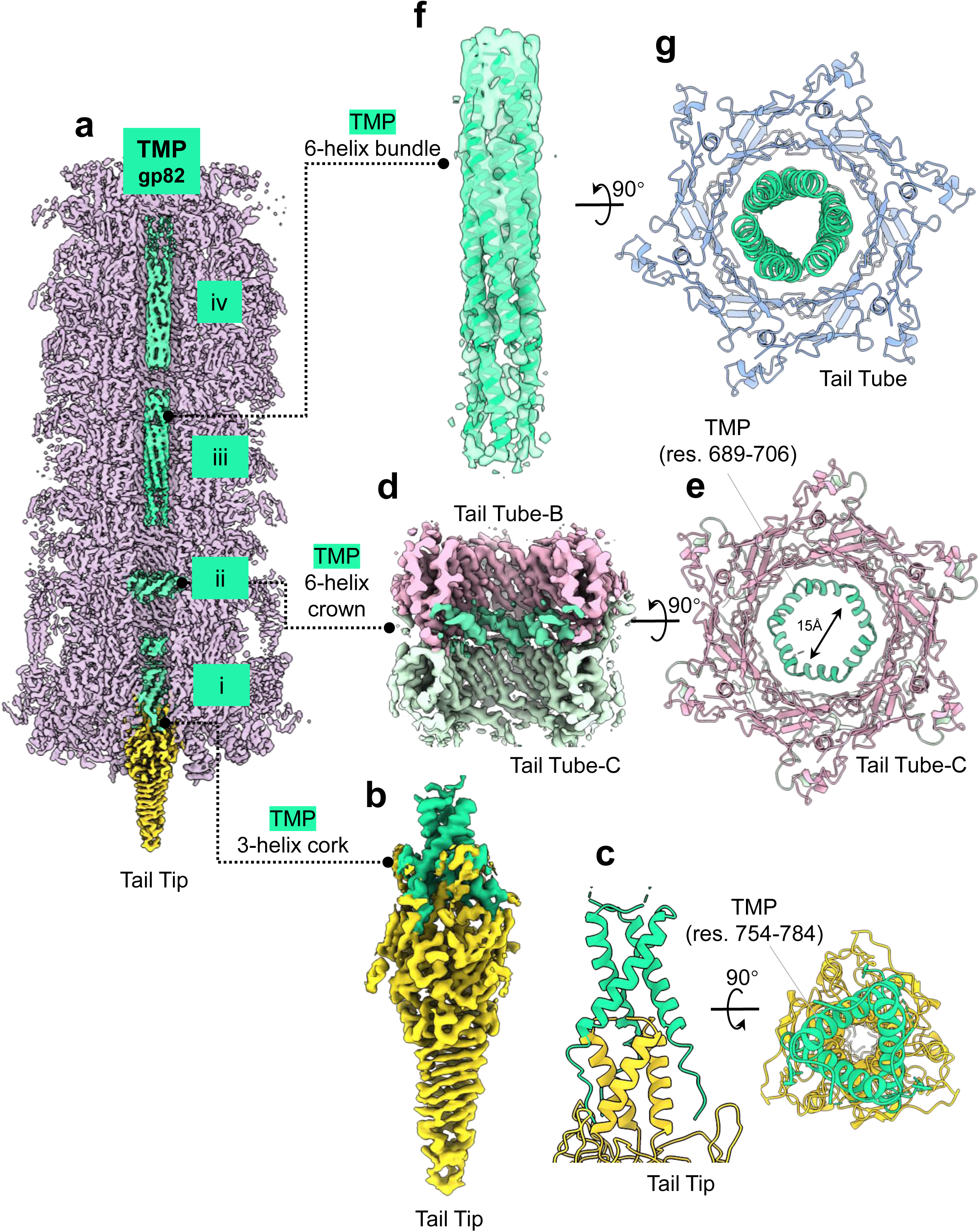
P7-1 Tape Measure Protein. (a) Cut-out view of P7-1 extended tail from a C3 density map calculated at a resolution of 4.5 Å. The extended tail and tail tip are colored gray and yellow, while the density for TMP is green. (b) Magnified view of the C3 density of the TMP 3-helix cork (green) bound to the N-terminal coiled-coil of the tail tip. (c) A ribbon diagram of the tip:TMP (res. 754-784) interaction is shown in side and top views. (d) A magnified view of the C3 density for the TMP 6-helix crown (green) located at the interface between the tail tube-B (pink) and tail tube-C (light green) hexamers of the cap subcomplex. (e) A ribbon diagram of the tail lumen, filled with six copies of the TMP (res. 689-706), is presented as a cut-out view. (f) A magnified view of the C3 density for the TMP 6-helix bundle filling the tail lumen. A polyalanine model for six helices was placed inside the density to help visualize the presence of six α-helices, each comprising approximately 70 residues. (g) A ribbon diagram of the tail lumen, filled with six helical densities of TMP, is presented in a cut-out view.

The second TMP region visible in the C3 reconstruction consists of a helical hexamer (the TMP 6-helix crown) (Fig. 9a(ii), d), formed by three α-helical hairpins made up of TMP residues 689-706. This ring is located at the inner interface between Tail Tube-B and Tail Tube-C (Fig. 9d), marking the first nonsymmetric interface in the repetitive tail tube lumen. TMP 6-helix crown inserts into the acidic tail tube interface, primarily bonding through Arg 691 and limiting the tail tube lumen to about 15 Å (Fig. 9e).

Finally, extensive elongated density for TMP fills the entire tail tube, which in the C3 map appears to form two 6-helix bundles. The bottom bundle (labeled as (iii) in Fig. 9a) is more continuous. It can be tentatively traced with six poly-alanine α-helices of approximately 70 residues each (Fig. 9f). The top bundle (labeled as (iv) in Fig. 9a) is less continuous and tapers off toward the top of the tail lumen, where the thinner density suggests that TMP lost a helical fold.

Since the cryo-EM reconstruction does not directly reveal whether TMP is trimeric or hexameric inside the P7-1 tail, we examined whether the tail volume can hold six copies of the TMP. To answer this question, we used the Matthews coefficient probability, which has been used in protein crystallography for decades to estimate the approximate number of molecules in a crystallographic asymmetric unit ^37^. The P7-1 tail lumen has a diameter of approximately 35.5 Å, calculated from juxtapposed arginine side chains lining the channel, and a length of roughly 1,100 Å. This dimension can vary depending on where the DNA density ends and where the TMP begins, but we assume that TMP begins at the start of the sheath (Fig. 3b). Thus, the volume of this cylinder is V = h x r^2^ x π = 1,100 x 315.06 x 3.14 = 1,088,217.2 A^3^. We then approximated the P7-1 tail lumen as a triclinic unit cell (containing only one asymmetric unit) with dimensions a = 31.4 Å, b = 31.4 Å, and c = 1,100 Å, resulting in a unit volume of 1,084,556 Å³, which is very similar to the volume of the cylinder described above. Assuming six copies of TMP (M.W. 85.9 kDa), the Matthews coefficient VM (V/M.W.) is 2.1 Å³/Da, which is a plausible value for a hydrated unit cell since it falls within the typical range of 1.62 to 3.53 Å³/Da. This corresponds to a unit cell with approximately 41.6% solvent content, similar to the tail tube filled with TMP observed experimentally by cryo-EM (Fig. 9g). Therefore, it is physically reasonable to have six copies of TMP inside the P7-1 tail lumen, supported by the presence of six identical helices in the TMP 6-helix crown (Fig. 9d). In contrast, three copies of TMP, as suggested by TMP 3-helix cork (Fig. 9b,c), lead to an unlikely Matthews coefficient of 4.2 Å³/Da and an estimated 70.8% solvent content, which may be too high to observe ordered TMP density inside P7-1 tail through single-particle analysis.

## DISCUSSION

*Pakpunaviruses* are of significant biomedical interest due to their potent bactericidal activity and broad host specificity ^38^. More than 60 *Pakpunavirus* genomes have been sequenced to date, and this number is expected to grow as large-scale sequencing efforts expand. However, the absence of structural and functional information has limited accurate annotation of their proteins. Here, using cryo-EM, localized reconstruction, proteomics, and bioinformatics, we annotated 19 structural genes of the *Pseudomonas* phage P7-1 and captured the virion in both extended, DNA-filled, and contracted, DNA-empty conformations. This structural atlas reveals several unanticipated features of P7-1 biology.

First, P7-1 virions exhibit remarkable stability. More than 95% of vitrified virions retained extended tails three months after purification, in contrast to the *Pbunavirus* E217, which shows ∼30% contracted, empty particles within days ^14^. P7-1 was also resistant to alkalinization, which triggered contraction in E217, while only prolonged acidification induced contraction in P7-1, and even then, most particles retained DNA. Fewer than 3% of contracted particles released their genome, confirming that contraction is necessary but not sufficient for genome ejection ^39^. Our cryo-EM reconstruction suggests that stability arises from two distinct tail fibers, STF and LTF, reminiscent of bacteriophage Sf14 ^40^. STFs are tightly integrated into the sheath lattice, where their bonding stabilizes the helical Triplex bundle and multiple sheath layers, physically blocking premature contraction ^30^. In contrast, LTFs are more loosely positioned, likely acting as flexible probes for host recognition. Together, these dual fibers provide both mechanical stability and infection specificity.

Second, P7-1 encodes two tail fibers per Triplex complex, yielding 12 fibers per baseplate. This arrangement arises from a duplication of the fiber attachment loop (FAL) in the Triplex small subunit gp89, a feature absent in *Pbunaviruses*. In this respect, P7-1 resembles phage T4, which also carries short and long fibers ^41,42^. Like T4, the larger genome of *Pakpunaviruses* (∼30% larger than most *Pbunaviruses*) is associated with greater fiber complexity, although the baseplate remains comparable in size to *Pbunaviruses* and smaller than that of T4. Functional data indicate that replacing P7-1 LTF with heterologous LTFs from other *Pakpunaviruses* alters host range, supporting a role for LTFs in recognizing strain-specific O-antigens of *P. aeruginosa* ^14^. STFs likely serve as secondary anchors via conserved receptors such as core LPS. Nonetheless, significant host-range overlap among phages with divergent LTFs suggests that multiple attachment mechanisms contribute to infection. Ongoing genetic work will clarify the relative roles of tail fibers and baseplate proteins in host specificity.

Third, we identified an oligomerization mismatch in the Tape Measure Protein (TMP). At the distal tail tip, TMP adopts three-fold symmetry, while within the lumen it exhibits six-fold symmetry, as reported for other Myoviruses ^14,43^. Six ordered helices (residues 689–706) were resolved, confirming that TMP exists in six copies within the extended tail. A theoretical calculation based on the inner tail volume confirmed this stoichiometry, showing that six copies of an 85.9 kDa TMP are chemically compatible with the volume of the P7-1 tail. The observed TMP structure visualized within the extended P7-1 tail likely represents a pre-ejection conformation, consistent with evidence that TMP refolds during genome delivery ^44^. TMP is multifunctional: in addition to determining tail length during assembly, it may facilitate genome ejection by forming a channel across the host membrane ^19,45^. Such roles are well established in *Siphoviridae* and supported by studies of λ ^46,47^, HK97 ^36^, and T5 ^48^. Moreover, TMP can encode peptidoglycan hydrolase domains ^49^. In P7-1, we identified a putative Cpl-7 lysozyme-like domain (residues 641–674), though its activity remains to be confirmed biochemically. Further structural and functional studies are needed to define TMP rearrangement after contraction and its precise role in genome ejection.

In summary, this work establishes P7-1 as the first *Pakpunavirus* phage with a structurally annotated virion from head to baseplate. By integrating cryo-EM with proteomics and comparative genomics, we provide insights into virion stability, fiber-mediated host recognition, and the potential role of TMP in genome delivery. This 3D atlas lays the foundation for mapping functional mutations and provides principles for the rational engineering of therapeutic phages with enhanced stability, specificity, and efficacy.

## METHODS

### Origin and characteristics of *Pseudomonas* phage P7-1

P7-1 was isolated from sewer samples collected in Southern California. Fermentation and purification were accomplished using proprietary methods to achieve clinical levels of purity and a titer of 1 × 10^13^ PFU/mL. P7-1 was sequenced on an Illumina MiSEQ and was assembled and annotated at Armata Pharmaceuticals using proprietary pipelines. The resulting genome is 93,745 bp long and contains 1,143 base-long Direct Terminal Repeats. The coding sequences relevant to this work are published in Supplementary Table 1. Bioinformatics analyses were performed using the most up-to-date version of Geneious Prime 2025.2 (https://www.geneious.com). Genomes of publicly available *Pakpunaviruses* were downloaded from NCBI, and their accession numbers are listed in Supplementary Table 3. Homologs of structural proteins for all phages were identified from whole genome alignments and corrected as necessary to match P7-1’s coding sequences. Genomes that lacked annotations were first annotated using the same pipelines as those used for the assembly and annotation of P7-1.

### Vitrification and data collection

3.0 µL of P7-1 virions, measured at a PFU of 1 x 10^13^ phages/mL, was applied to a 200-mesh copper Quantifoil R 2/1 holey carbon grid (EMS) previously glow-discharged for 60 sec at 15 mA using an easiGlow (PELCO). The grid was blotted for 7.5 sec at blot force 2 and vitrified immediately in liquid ethane using a Vitrobot Mark IV (Thermo Scientific). Cryo-grids were screened on a 200 kV Glacios (Thermo Scientific) equipped with a Falcon4 detector (Thermo Scientific) at Thomas Jefferson University. EPU software (Thermo Scientific) was used for data screening using the accurate positioning mode. For high-resolution data collection of P7-1 with an extended tail, micrographs were collected on a Titan Krios (Thermo Scientific) microscope operated at 300 kV and equipped with a K3 direct electron detector camera (Gatan) at the National Cryo-Electron Microscopy Facility (NCEF) at the Frederick National Laboratory, MD. To induce P7-1 tail contraction, virions were incubated at pH 4.5 for 2 hours. A dataset was collected at Stanford-SLAC CryoEM Center (S^2^C^2^) on a 300 kV Krios equipped with a Falcon 4 direct detector.

### Liquid chromatography/mass spectrometry (LC/MS/MS) analysis

Phage samples were treated with 12 mM sodium lauryl sarcosine, 0.5% sodium deoxycholate, and 50 mM triethyl ammonium bicarbonate (TEAB), heated to 95 °C for 10 min, and then sonicated for 10 min, followed by the addition 5 mM tris(2-carboxyethyl) phosphine and 10 mM chloroacetamide to entirely reduce and alkylate the proteins in the sample. The sample was then subjected to trypsin digestion overnight (1:100 w/w trypsin added two times). Following digestion, the sample was acidified, lyophilized, and then desalted before injection onto a laser-pulled nanobore C18 column with 1.8 μm beads. This was followed by ionization through a hybrid quadrupole-Orbitrap mass spectrometer. The most abundant proteins were identified by searching the experimental data against a phage protein database, a *Pseudomonas* host protein database, and a common contaminant database using the MASCOT algorithm ^50^.

### Cryo-EM SPA

All micrographs of phage P7-1 were motion-corrected with MotionCorr2 ^51^. RELION’s implementation of motion correction was applied to the micrographs with options of dose-weighted averaged micrographs and the sum of non-dose weighted power spectra every 4 e^-^/Å^2^. Contrast Transfer Function (CTF) was estimated using CTFFIND4 ^52^. After initial reference picking and 2D classification, particles were subjected to a reference-free low-resolution reconstruction without imposing symmetry. The particles were then 3D classified into four classes, with I4 symmetry imposed. Among the four classes, the best one was selected and underwent 3D auto-refinement to finely align the particles. The particles were then expanded according to I4 symmetry using RELION’s *relion_particle_symmetry_expand* function to obtain 60 times the initial particles. A cylindrical mask (r = 200 Å) was generated using SCIPION 3.0 ^53^ and then resampled onto a reference map covering the five-fold vertex in Chimera ^54^. The cylindrical mask was then used for non-sampling 3D classification (as implemented in RELION 3.1.2 ^55,56^) without imposing symmetry to search for the tail. Locally aligned particles were then combined, and duplicate particles were removed. The initial localized reference map was reconstructed directly from one of the classes using RELION’s *ab initio* 3D Initial Model. Selected 3D classes were auto-refined using C5 symmetry, followed by five-fold particle expansion. The expanded particles underwent a third 3D classification, and the map was symmetrized using C12 and C6 symmetries, which yielded the best densities for the portal/head-to-tail and collar/gateway/tube/sheath proteins, respectively.

To reconstruct the baseplate, we manually picked 10,400 particles containing the distal tip of the P7-1 tail and reconstructed it with C3 symmetry, as C6 symmetry would result in a loss of density features. For the contracted sheath, P7-1 with contracted tails were picked from 7,832 movies collected on a 300 kV Krios/Falcon 4 and processed as detailed above. After extensive particle picking and 2D classification, 10,329 contracted tail particles were reconstructed in C6 symmetry. All steps of SPA, including 2D Classification, 3D classification, 3D refinement, CTF refinement, particle polishing, post-processing, and local resolution calculation, were carried out using RELION 3.1.2 ^55,56^. The final densities were sharpened using *phenix.autosharpen* ^57^. RELION_*postprocess* ^55,56^ was used for local resolution estimation. All cryo-EM data collection statistics are in Table 1.

### *De novo* model building, oligomer generation, and refinement

All *de novo* atomic models presented in this paper were built using Coot ^58^ and ChimeraX ^59^. We used four different maps for model building: (***i***) a 3.3 Å C5-averaged localized reconstruction ^16^ of the mature capsid (Supplementary Fig. 2a) yielded models of the capsid (gp71) and the decoration protein (gp70); (***ii***) 3.6 Å C12- and C6-averaged localized reconstructions of the extended neck and tail (Supplementary Fig. 2b) were used to build the dodecameric portal (gp67) and HT-adaptor (gp72), as well as hexameric collar (gp74), gateway (gp75), tail sheath (gp76), and tail tube (gp77); (***iii***) a 3.6 Å C3-averaged reconstruction of the extended baseplate (Supplementary Fig. 2c) yielded models of the tube-B (gp78), tube-C (gp79), ripcord (gp83), adaptor-1 (gp84), hub (gp85), tip (gp86), adaptor-2 (gp87), Triplex subunits (gp88-a/gp88-b/gp89). ModelAngelo ^35^ was used to *de novo* build TMP (gp82) residues 689-706 and 754-784. The same C3 reconstruction of the extended baseplate and tail bottom, but calculated at 4.2 Å resolution, was used to dock AlphaFold3 ^33^ models of STF and LTF. Both fibers were fit into the density using the ‘fit-into-map’ command in ChimeraX ^59^. (***iv***) A 4.2 Å C6-averaged reconstruction of the contracted baseplate (Supplementary Fig. 2d) was used to dock all the factors observed in the contracted tail and baseplate subunits. All atomic models, except for LTF, were refined against the different densities described above through multiple rounds of rigid-body, real-space, and B-factor refinement with *phenix.real_space_refinement* ^60^. Figure 2 shows all final models generated in this study (except for TMP fragments), and Table 1 presents the relative map and model statistics.

### Structural analysis

All ribbon and surface representations were generated using ChimeraX ^59^. Drawings of electron density maps and local resolution maps were generated using ChimeraX ^59^. Structural neighbors and flexible regions were identified using Foldseek ^61^. RMSD between superimposed PDBs was calculated using SuperPose Version 1.0 (superpose.wishartlab.com) ^62^. Binding interfaces were analyzed using PISA ^63^ and PDBsum ^64^ to determine bonding interactions, interatomic distances, and bond types. Structure prediction was done using AlphaFold ^17^, and the hexameric TMP was generated using AlphaFold3 ^33^.

### Statistics and reproducibility

The cryo-EM data were collected from several grids. Micrographs with poor ice were excluded from the reconstruction pool by visual inspection. Data collection, processing, and refinement statistics are summarized in Table 1. All atomic models were validated using MolProbity ^65^. All RMSDs in the Cα position between superimposed structures were calculated using Coot ^58^.

## ACKNOWLEDGMENTS

We thank the staff at SLAC and NCEF for assistance in remote data collection. This work was supported by National Institutes of Health grants R01 GM100888 and R35 GM140733. Electron microscopy was carried out in the UAB Cryo-EM Facility (RRID:SCR_025450), supported by the Institutional Research Core Program and O’Neal Comprehensive Cancer Center (NIH grant P30 CA013148), with additional funding from NIH grant S10 OD024978. We thank Dr. James Kizziah for assisting with the in-house data collection. A portion of this work was carried out at NCEF (supported by contract 75N91019D00024) and Stanford-SLAC CryoEM Center (S^2^C^2^), which is supported by the NIH Common Fund Transformative High-Resolution Cryo-Electron Microscopy program (U24 GM129541).

## AUTHOR CONTRIBUTIONS STATEMENT

F.L., N.B., C-F.D.H., R.Y., and G.C. performed all steps of the cryo-EM data collection and analysis, deposition of atomic coordinates, and maps. R.K.L. helped with biochemical methods. G.C., P.K., D.B., and S.L. supervised the entire project. F.L., S.L., and G.C. wrote the paper. Z.K., R.G., A.S., S.B., and S.L. isolated, amplified, and purified P7-1 for cryo-EM analysis. A.S. analyzed the LC/MS MS data in conjunction with the service provider. R.G. and S.L. sequenced and analyzed the genome of P7-1. All authors contributed to the writing and editing of the manuscript.

## DATA AVAILABILITY

The atomic models and three-dimensional reconstructions described in this paper are available in the Protein Data Bank (9ZBI, 9ZBS, 9ZD8, 9ZD7, 9ZDK) and Electron Microscopy Data Bank (EMD-73989, EMD-73999, EMD-74057, EMD-74056, EMD-74065), respectively. All other data are available from the corresponding author upon reasonable request.

