## Supplemental Figs 1-5 and Tables 1-5 for "Structural atlas of *Pakpunavirus* P7-1 reveals determinants of virion stability and genome ejection"

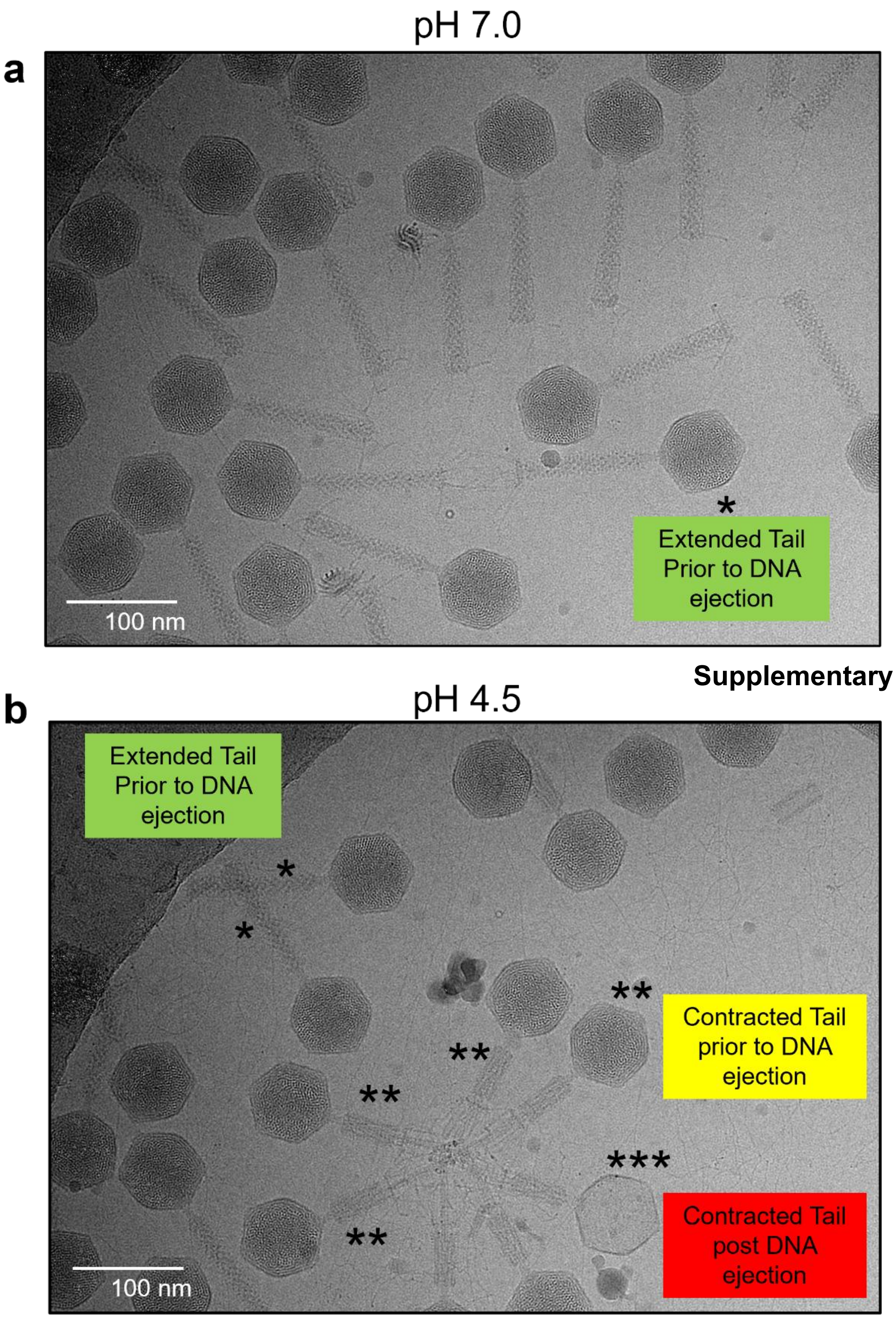


**Supplementary Figure 1. Cryo-EM micrographs.** (**a**) P7-1 vitrified at neutral pH 7.0. (**b**) P7-1 vitrified at acidic pH 4.5. Three types of particles are visible, indicated by (*), (**) and (***).

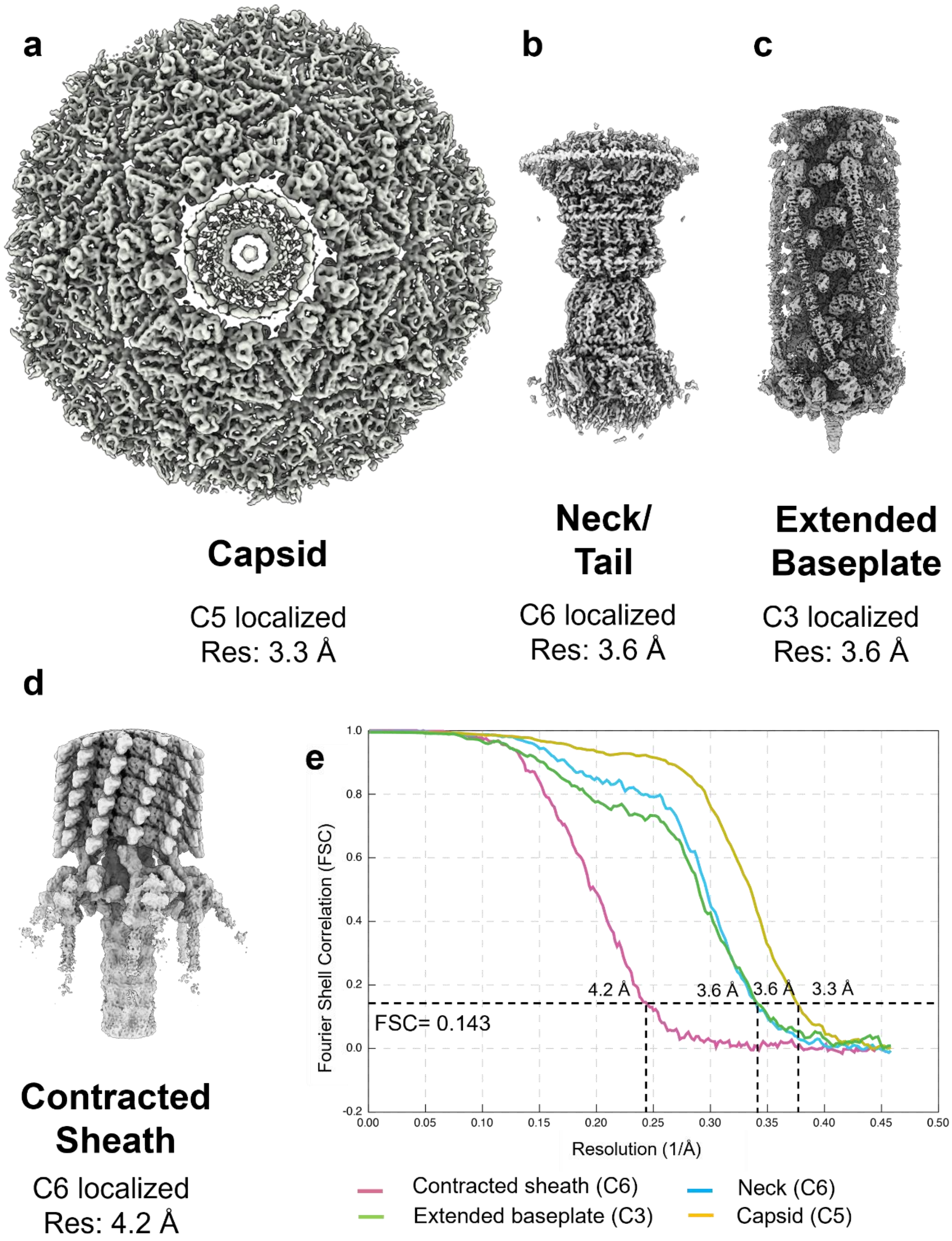


**Supplementary Figure 2. Cryo-EM reconstructions.** (**a**) Cryo-EM reconstructions determined at pH 7.0 in this study and used to build atomic models are shown in Figure 2. (**b**) Cryo-EM reconstruction of the contracted sheath determined at pH 4.5. (**c**) Fourier Shell Correlation (FSC) curves for all cryo-EM reconstructions in this study (e.g., the resolution is indicated at the 0.143 cut-off).

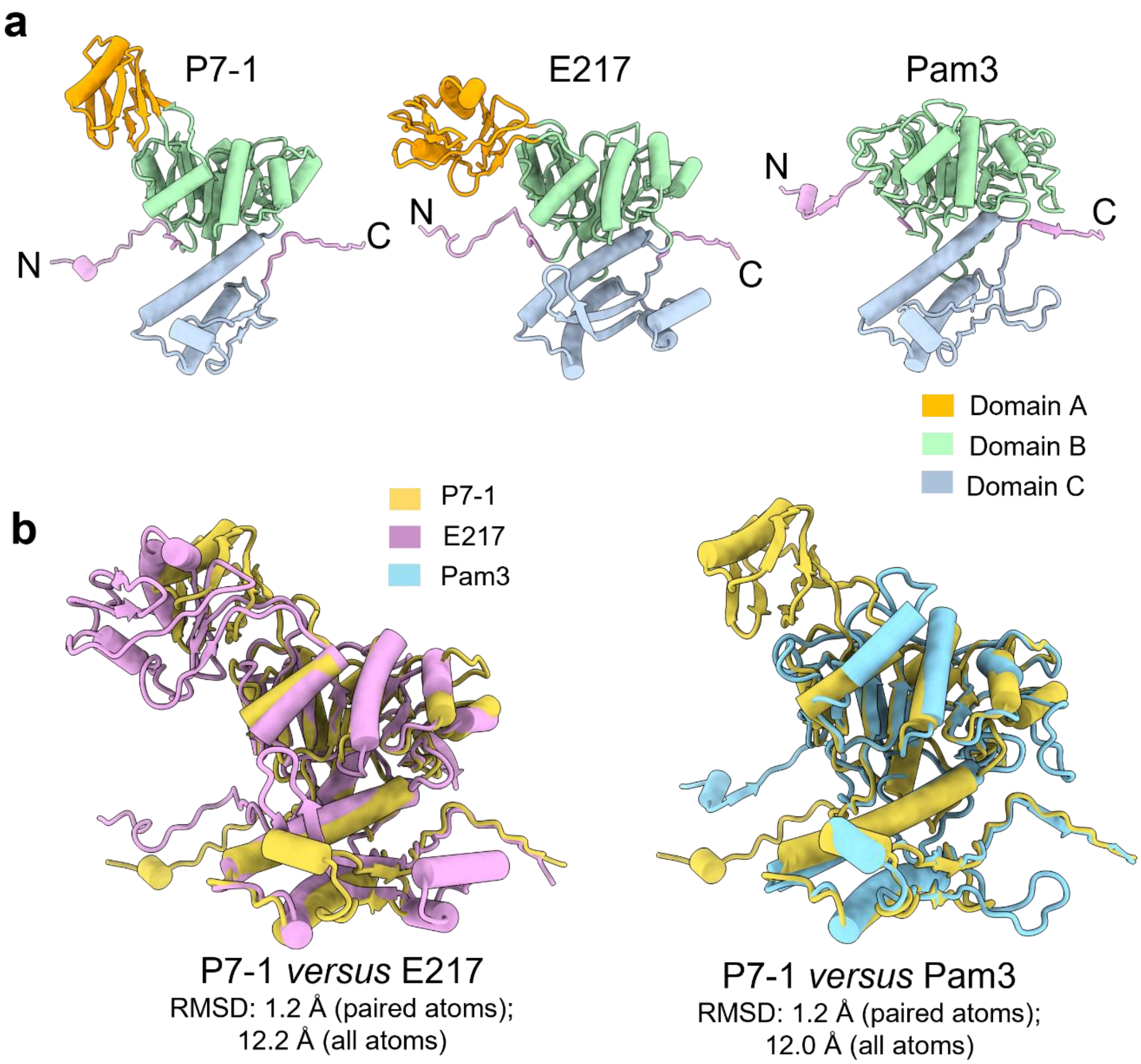


**Supplementary Figure 3. Structure and conservation of the sheath proteins of phages P7-1, E217, and Pam3.** (**a**) Three-dimensional structure of the sheath proteins from phages P7-1 (gp76), E217 (gp31), and Pam3 (gp13), with domains color-coded consistently. (**b**) Structural superimposition of the P7-1 sheath protein (orange) with the counterparts from E217 (purple) and phage Pam3 (cyan).

| | gp67 Portal Protein | gp70 Head Decoration Protein | gp71 Major Capsid Protein | gp72 Head-To-Tail Protein | gp74 Collar Protein | gp75 Gateway Protein | gp76 Tail Sheath Protein | gp77 Tail Tube Protein | gp78 Tail Tube Protein B | gp79 Tail Tube Protein C | gp82 Tape Measure Protein | gp83 Ripcord Protein | gp84 Adaptor Protein 1 | gp85 Baseplate Hub Protein | gp86 Tail Tip Protein | gp87 Adaptor Protein 2 | gp88 Large Baseplate Wedge Protein | gp89 Small Baseplate Wedge Protein | gp90 Long Tail Fiber Protein | gp92 Short Tail Fiber Protein |
|---|---|---|---|---|---|---|---|---|---|---|---|---|---|---|---|---|---|---|---|---|
| vB-PaeM-YQ78 | 99.7 | 100.0 | 100.0 | 98.7 | 100.0 | 97.3 | 98.4 | 100.0 | 99.4 | 100.0 | 99.1 | 100.0 | 100.0 | 100.0 | 100.0 | 100.0 | 99.6 | 97.2 | 100.0 | 99.6 |
| HJ01 | 99.9 | 99.3 | 99.7 | 98.7 | 100.0 | 100.0 | 98.6 | 100.0 | 100.0 | 100.0 | 99.1 | 100.0 | 100.0 | 100.0 | 100.0 | 99.2 | 99.4 | 99.6 | 99.9 | 99.6 |
| vB_VIPPAEUMC01 | 99.5 | 100.0 | 100.0 | 100.0 | 100.0 | 100.0 | 98.6 | 100.0 | 100.0 | 100.0 | 99.0 | 100.0 | 100.0 | 100.0 | 100.0 | 100.0 | 99.8 | 99.6 | 99.7 | 99.8 |
| PA45_GUMS | 99.4 | 100.0 | 100.0 | 100.0 | 100.0 | 99.5 | 98.8 | 100.0 | 100.0 | 100.0 | 100.0 | 98.4 | 100.0 | 100.0 | 100.0 | 100.0 | 99.6 | 99.6 | 99.4 | 99.4 |
| pPA-3099-2aT.2 | 99.1 | 100.0 | 100.0 | 98.7 | 100.0 | 100.0 | 98.4 | 100.0 | 100.0 | 100.0 | 100.0 | 98.4 | 100.0 | 99.7 | 100.0 | 100.0 | 99.4 | 99.6 | 99.4 | 99.2 |
| vB_PaeM_B31 | 99.7 | 100.0 | 100.0 | 98.7 | 100.0 | 100.0 | 98.6 | 100.0 | 100.0 | 100.0 | 99.0 | 100.0 | 99.2 | 100.0 | 100.0 | 100.0 | 99.4 | 99.6 | 99.1 | 99.2 |
| vB_PaeM_C2-10_Ab02 | 99.5 | 100.0 | 100.0 | 100.0 | 100.0 | 100.0 | 98.8 | 100.0 | 100.0 | 100.0 | 100.0 | 98.4 | 100.0 | 100.0 | 100.0 | 100.0 | 99.6 | 99.6 | 99.1 | 99.4 |
| vB_Paer_Ps12 | 99.2 | 99.3 | 100.0 | 100.0 | 100.0 | 100.0 | 99.5 | 100.0 | 100.0 | 100.0 | 99.5 | 100.0 | 99.2 | 99.7 | 100.0 | 100.0 | 99.6 | 99.2 | 99.0 | 99.4 |
| PaP1 | 98.8 | 99.3 | 100.0 | 100.0 | 100.0 | 100.0 | 99.5 | 100.0 | 96.4 | 100.0 | 99.1 | 100.0 | 99.2 | 99.7 | 100.0 | 100.0 | 99.2 | 100.0 | 96.9 | 99.6 |
| vB_Paer_Ps25 | 99.2 | 99.3 | 100.0 | 100.0 | 100.0 | 100.0 | 99.8 | 100.0 | 100.0 | 100.0 | 99.5 | 100.0 | 99.2 | 99.7 | 100.0 | 100.0 | 99.6 | 100.0 | 98.8 | 99.4 |
| vB_PaeM_C2-10_Ab08 | 99.1 | 100.0 | 100.0 | 98.1 | 100.0 | 100.0 | 99.8 | 100.0 | 100.0 | 100.0 | 100.0 | 97.6 | 100.0 | 100.0 | 100.0 | 100.0 | 99.6 | 96.8 | 98.8 | 99.4 |
| vB_PaeM_C2-10_Ab10 | 99.1 | 100.0 | 100.0 | 98.7 | 98.4 | 97.3 | 99.3 | 100.0 | 100.0 | 100.0 | 100.0 | 97.6 | 100.0 | 100.0 | 100.0 | 100.0 | 99.6 | 96.8 | 98.4 | 99.4 |
| vB_PaeM_LCK69 | 99.5 | 100.0 | 100.0 | 98.1 | 100.0 | 98.9 | 98.6 | 100.0 | 100.0 | 100.0 | 99.9 | 100.0 | 99.2 | 99.7 | 100.0 | 100.0 | 99.2 | 100.0 | 98.6 / 98.3 | 99.2 |
| vB_PaeM_C2-10_Ab1 | 99.1 | 100.0 | 100.0 | 98.7 | 98.4 | 97.3 | 98.6 | 100.0 | 100.0 | 100.0 | 100.0 | 97.2 | 100.0 | 100.0 | 100.0 | 100.0 | 99.6 | 99.6 | 98.4 | 99.2 |
| vB_Paer_PsIn | 96.7 | 100.0 | 100.0 | 100.0 | 100.0 | 100.0 | 99.8 | 100.0 | 100.0 | 100.0 | 99.1 | 99.2 | 100.0 | 100.0 | 100.0 | 100.0 | 99.6 | 100.0 | 98.4 | 99.4 |
| Ab15 | 96.7 | 99.3 | 100.0 | 100.0 | 100.0 | 100.0 | 98.8 | 100.0 | 100.0 | 100.0 | 100.0 | 99.6 | 100.0 | 100.0 | 100.0 | 100.0 | 99.6 | 99.2 | 98.4 | 99.4 |
| PIAS | 99.9 | 100.0 | 100.0 | 100.0 | 100.0 | 100.0 | 98.1 | 100.0 | 99.4 | 100.0 | 99.0 | 100.0 | 99.2 | 100.0 | 100.0 | 100.0 | 99.6 | 99.2 | 98.2 | 99.4 |
| vB_Paer_PsCh | 99.2 | 99.3 | 100.0 | 100.0 | 100.0 | 100.0 | 99.8 | 100.0 | 100.0 | 100.0 | 99.1 | 99.2 | 100.0 | 99.3 | 100.0 | 100.0 | 99.6 | 100.0 | 98.2 | 99.0 |
| vB_PaeS_B8 | 99.5 | 100.0 | 100.0 | 98.7 | 100.0 | 97.3 | 99.1 | 100.0 | 100.0 | 100.0 | 99.0 | 98.4 | 100.0 | 99.7 | 100.0 | 100.0 | 99.6 | 99.6 | 98.2 | 99.2 |
| YS35 | 99.7 | 100.0 | 100.0 | 99.4 | 100.0 | 97.3 | 99.1 | 100.0 | 100.0 | 100.0 | 99.0 | 98.4 | 100.0 | 100.0 | 100.0 | 100.0 | 99.6 | 99.2 | 98.2 | 99.2 |
| GEC_PNG3 | 99.0 | 100.0 | 100.0 | 100.0 | 100.0 | 100.0 | 99.5 | 100.0 | 100.0 | 100.0 | 99.9 | 100.0 | 99.2 | 99.7 | 100.0 | 100.0 | 99.4 | 99.6 | 98.1 | 99.4 |
| GEC_MRC | 99.1 | 99.3 | 99.7 | 99.4 | 94.4 | 95.7 | 99.1 | 100.0 | 100.0 | 99.4 | 100.0 | 99.6 | 100.0 | 100.0 | 100.0 | 100.0 | 99.6 | 99.6 | 97.8 | 99.4 |
| PA10 | 99.9 | 99.3 | 99.7 | 99.4 | 100.0 | 95.7 | 99.1 | 100.0 | 100.0 | 100.0 | 99.1 | 100.0 | 99.2 | 99.7 | 100.0 | 99.2 | 99.2 | 100.0 | 97.8 | 99.6 |
| PAK_P2 | 99.5 | 100.0 | 100.0 | 100.0 | 100.0 | 100.0 | 99.8 | 100.0 | 100.0 | 100.0 | 99.1 | 100.0 | 94.9 | 98.4 | 100.0 | 100.0 | 99.4 | 100.0 | 97.5 | 99.2 |
| PAK_P4 | 99.9 | 100.0 | 100.0 | 100.0 | 100.0 | 97.3 | 98.6 | 100.0 | 100.0 | 100.0 | 99.1 | 100.0 | 99.2 | 99.7 | 100.0 | 100.0 | 99.4 | 100.0 | 96.7 | 99.6 |
| K5 | 98.8 | 99.3 | 100.0 | 100.0 | 100.0 | 100.0 | 98.6 | 100.0 | 100.0 | 100.0 | 99.1 | 100.0 | 99.2 | 99.7 | 100.0 | 100.0 | 99.2 | 100.0 | 96.7 | 99.4 |
| K8 | 98.8 | 99.3 | 100.0 | 99.4 | 100.0 | 100.0 | 98.6 | 100.0 | 100.0 | 100.0 | 99.1 | 100.0 | 99.2 | 99.7 | 100.0 | 100.0 | 99.2 | 100.0 | 96.7 | 99.4 |
| phipa10 | 98.8 | 100.0 | 100.0 | 99.4 | 99.2 | 100.0 | 98.6 | 100.0 | 100.0 | 100.0 | 99.8 | 100.0 | 99.2 | 99.7 | 100.0 | 100.0 | 99.4 | 99.6 | 96.6 | 99.2 |
| PJNP029 | 98.8 | 99.3 | 100.0 | 100.0 | 100.0 | 100.0 | 98.6 | 100.0 | 100.0 | 100.0 | 99.0 | 100.0 | 99.2 | 99.7 | 100.0 | 100.0 | 99.2 | 100.0 | 96.0 | 99.2 |
| vB_PaeM_B55 | 99.2 | 100.0 | 100.0 | 99.4 | 100.0 | 100.0 | 99.1 | 100.0 | 100.0 | 100.0 | 99.0 | 100.0 | 99.2 | 99.7 | 100.0 | 100.0 | 99.2 | 100.0 | 96.0 | 99.2 |
| ITTPL | 96.6 | 99.3 | 99.7 | 100.0 | 97.6 | 96.8 | 96.3 | 100.0 | 99.4 | 91.8 | 91.8 | 94.9 | 95.8 | 96.4 | 98.4 | 96.8 | 94.1 | 96.7 | 87.8 | 95.4 |
| PAK_P1 | 98.8 | 99.3 | 100.0 | 100.0 | 99.2 | 99.5 | 98.1 | 100.0 | 99.4 | 91.2 | 91.9 | 94.5 | 95.8 | 96.4 | 99.2 | 95.1 | 94.3 | 96.7 | 88.7 | 99.4 |
| vB_PaeM_MAG1 | 99.9 | 100.0 | 100.0 | 98.1 | 100.0 | 97.3 | 99.3 | 100.0 | 100.0 | 100.0 | 99.1 | 100.0 | 98.3 | 100.0 | 100.0 | 100.0 | 99.2 | 100.0 | 89.1 | 95.8 |
| EM | 85.2 | 97.1 | 98.3 | 98.7 | 93.7 | 96.3 | 95.3 | 100.0 | 98.8 | 95.6 | 93.2 | 97.2 | 95.7 | 97.4 | 92.3 | 95.9 | 95.5 | 95.9 | 73.2 | 92.4 |
| vB_PaM_EPA1 | 99.6 | 100.0 | 100.0 | 98.1 | 100.0 | 97.3 | 99.1 | 100.0 | 100.0 | 100.0 | 100.0 | 98.4 | 100.0 | 99.7 | 100.0 | 100.0 | 99.6 | 99.6 | 68.6 | 99.4 |
| Pa-U | 99.9 | 100.0 | 100.0 | 100.0 | 100.0 | 99.5 | 99.1 | 100.0 | 100.0 | 100.0 | 97.9 / 97.4 | 100.0 | 100.0 | 99.7 | 100.0 | 100.0 | 99.4 | 99.6 | 68.6 | 99.4 |
| SCUT-S2 | 99.4 | 98.5 | 100.0 | 98.7 | 100.0 | 97.3 | 99.1 | 100.0 | 100.0 | 100.0 | 100.0 | 98.4 | 100.0 | 99.7 | 100.0 | 100.0 | 99.6 | 99.6 | 68.6 | 99.2 |
| Zigelbrucke | 98.8 | 99.3 | 100.0 | 98.7 | 99.2 | 100.0 | 99.1 | 100.0 | 100.0 | 100.0 | 99.1 | 100.0 | 99.2 | 99.7 | 100.0 | 100.0 | 99.2 | 100.0 | 67.7 | 99.6 |
| phiMK | 99.1 | 100.0 | 100.0 | 99.4 | 98.4 | 96.8 | 98.6 | 100.0 | 100.0 | 100.0 | 100.0 | 97.6 | 96.6 | 96.4 | 98.4 | 96.8 | 96.9 | 100.0 | 67.4 | 97.2 |
| PhL_UNISO_PA-DSM_ph0034 | 98.8 | 100.0 | 100.0 | 98.7 | 100.0 | 99.5 | 99.5 | 100.0 | 100.0 | 98.7 | 98.5 | 100.0 | 98.3 | 100.0 | 100.0 | 100.0 | 99.8 | 100.0 | 67.2 | 99.4 |
| vFB297 | 96.8 | 99.3 | 99.7 | 99.4 | 94.4 | 95.7 | 99.3 | 100.0 | 100.0 | 100.0 | 99.1 | 99.6 | 99.2 | 99.7 | 100.0 | 100.0 | 99.4 | 100.0 | 66.7 | 95.6 |
| 20Sep416 | 99.1 | 100.0 | 100.0 | 98.7 | 100.0 | 97.9 | 99.1 | 100.0 | 100.0 | 100.0 | 100.0 | 97.2 | 100.0 | 100.0 | 100.0 | 100.0 | 99.4 | 100.0 | 66.6 | 99.6 |
| PaoP5 | 99.9 | 100.0 | 98.8 | 99.4 | 100.0 | 100.0 | 98.8 | 100.0 | 99.4 | 91.8 | 91.9 | 94.5 | 96.6 | 96.4 | 98.4 | 96.8 | 96.7 | 100.0 | 66.6 | 98.2 |
| 20Sep418 | 99.5 | 100.0 | 100.0 | 99.4 | 100.0 | 97.3 | 98.6 | 100.0 | 100.0 | 99.4 | 99.0 | 97.6 | 94.1 | 98.4 | 100.0 | 100.0 | 98.2 | 99.6 | 66.4 | 99.8 |
| LPS-5 | 99.5 | 100.0 | 99.7 | 99.4 | 100.0 | 97.3 | 98.6 | 100.0 | 100.0 | 99.4 | 98.9 | 96.8 | 94.9 | 98.4 | 100.0 | 100.0 | 99.2 | 81.7 / 100.0 | 66.4 | 99.6 |
| PaSz-1_45_92k | 99.5 | 99.3 | 99.7 | 99.4 | 100.0 | 95.2 | 98.6 | 100.0 | 100.0 | 99.4 | 99.1 | 99.2 | 94.1 | 98.4 | 100.0 | 100.0 | 98.2 | 100.0 | 66.4 | 99.8 |
| PaSzW-1 | 99.9 | 100.0 | 100.0 | 99.4 | 100.0 | 95.2 | 98.8 | 100.0 | 100.0 | 99.4 | 95.1 | 94.5 | 96.6 | 96.4 | 98.4 | 96.8 | 96.7 | 100.0 | 66.4 | 98.2 |
| PaYy-2 | 99.5 | 99.3 | 100.0 | 98.7 | 100.0 | 97.3 | 98.8 | 100.0 | 100.0 | 100.0 | 100.0 | 100.0 | 98.3 | 99.7 | 100.0 | 100.0 | 99.4 | 100.0 | 66.4 | 99.6 |
| PaZq-1 | 99.9 | 100.0 | 100.0 | 99.4 | 100.0 | 95.2 | 98.8 | 100.0 | 100.0 | 99.4 | 95.1 | 94.5 | 96.6 | 96.4 | 98.4 | 96.8 | 96.7 | 100.0 | 66.4 | 98.2 |
| PseuPha1 | 99.0 | 100.0 | 100.0 | 100.0 | 100.0 | 99.5 | 98.6 | 100.0 | 100.0 | 100.0 | 95.6 | 94.9 | 94.1 | 98.4 | 100.0 | 100.0 | 98.2 | 99.6 | 66.4 | 99.8 |
| 908-1 | 99.5 | 100.0 | 100.0 | 99.4 | 100.0 | 97.3 | 98.6 | 100.0 | 100.0 | 99.4 | 99.0 | 97.6 | 94.1 | 98.4 | 100.0 | 100.0 | 98.2 | 99.6 | 66.4 | 99.8 |
| Baskent_P3_3B | 99.5 | 100.0 | 100.0 | 99.4 | 100.0 | 97.3 | 98.6 | 100.0 | 100.0 | 99.4 | 99.3 / 98.7 | 98.4 | 100.0 | 100.0 | 100.0 | 100.0 | 99.2 | 100.0 | 66.4 | 99.6 |
| bmx-p3 | 99.5 | 100.0 | 100.0 | 99.4 | 100.0 | 97.3 | 98.6 | 100.0 | 100.0 | 99.4 | 99.0 | 97.6 | 94.1 | 98.4 | 100.0 | 100.0 | 98.2 | 99.6 | 66.4 | 99.8 |
| SPA05 | 98.8 | 98.5 | 99.7 | 99.4 | 100.0 | 100.0 | 99.1 | 100.0 | 100.0 | 100.0 | 98.9 | 94.5 | 96.6 | 96.4 | 98.4 | 95.9 | 96.9 | 99.6 | 66.4 | 99.6 |
| SRT6 | 98.6 | 99.3 | 100.0 | 98.1 | 100.0 | 97.3 | 99.1 | 100.0 | 100.0 | 100.0 | 100.0 | 99.6 | 99.2 | 99.7 | 100.0 | 100.0 | 99.4 | 100.0 | 66.3 | 99.6 |
| C11 | 98.8 | 99.3 | 100.0 | 100.0 | 100.0 | 99.5 | 99.3 | 100.0 | 100.0 | 100.0 | 99.1 | 95.3 | 95.8 | 96.4 | 98.0 | 96.8 | 96.9 | 100.0 | 66.3 | 99.6 |
| Henu5 | 99.5 | 100.0 | 100.0 | 100.0 / 67.1 | 100.0 | 99.5 | 99.1 | 100.0 | 100.0 | 100.0 | 99.9 | 100.0 | 99.2 | 99.7 | 100.0 | 100.0 | 99.2 | 100.0 | 66.3 | 99.4 / 93.2 |
| PaGz-1 | 98.8 | 99.3 | 99.4 | 100.0 | 98.4 | 99.5 | 98.6 | 100.0 | 99.4 | 91.8 | 91.6 | 94.9 | 96.6 | 96.4 | 98.0 | 96.8 | 96.7 | 99.6 | 66.3 | 97.0 |
| Kat | 99.5 | 100.0 | 99.7 | 99.4 | 100.0 | 95.2 | 98.4 | 100.0 | 100.0 | 100.0 | 98.9 | 96.1 | 94.1 | 98.7 | 100.0 | 100.0 | 98.2 | 100.0 | 66.1 | 99.2 |
| SPA01 | 96.9 | 97.8 | 99.7 | 100.0 | 98.4 | 99.5 | 98.8 | 100.0 | 100.0 | 100.0 | 94.5 | 94.5 | 95.8 | 99.0 | 100.0 | 100.0 | 98.2 | 99.6 | 66.1 | 95.8 |
| JG004 | 99.1 | 100.0 | 100.0 | 99.4 / 100.0 | 100.0 | 100.0 | 83.6 / 98.6 | 100.0 | 100.0 | 100.0 | 98.7 | 98.8 | 98.3 | 98.7 | 99.6 | 100.0 | 99.4 | 100.0 | 65.4 | 83.7 / 99.2 |

**Supplementary Figure 4. Comparison of the primary sequences of structural proteins from sequenced *Pakpunaviruses*.**

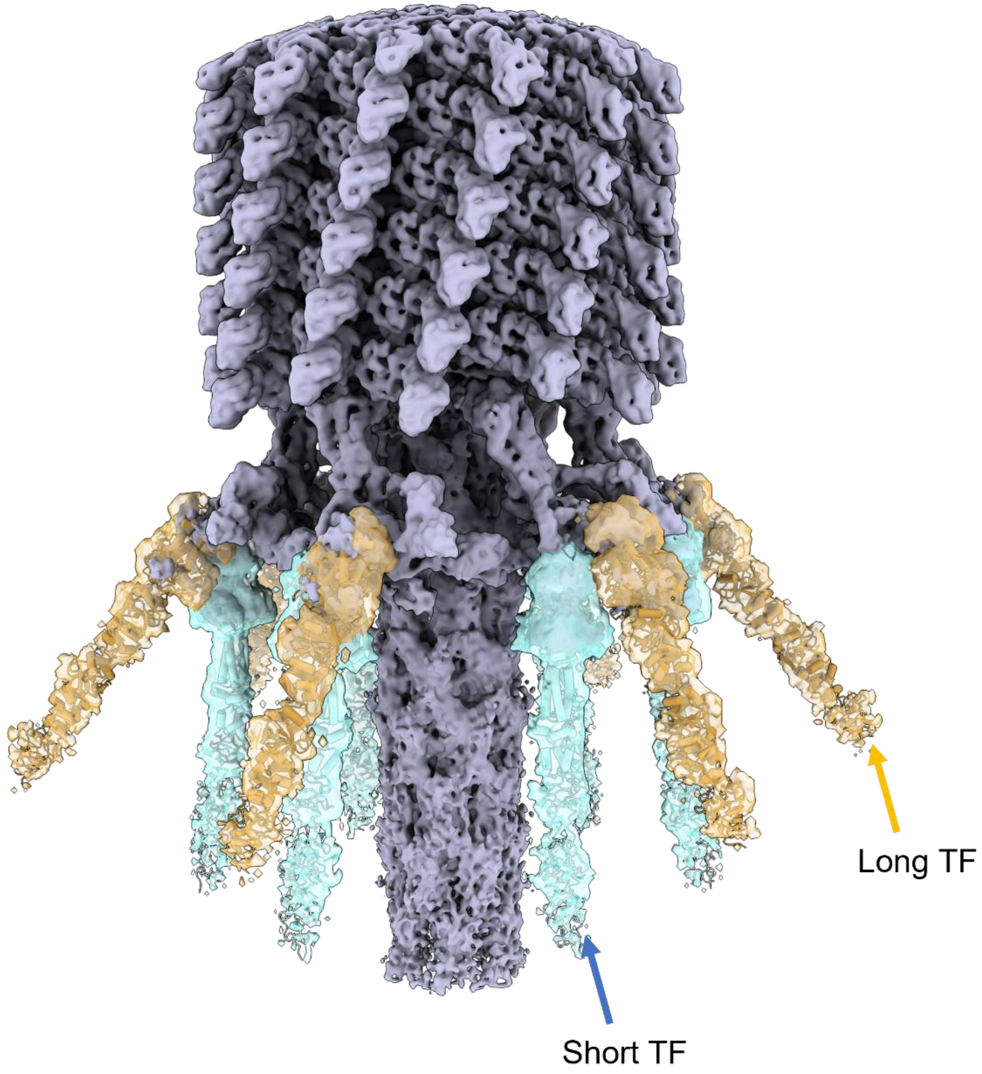


**Supplementary Figure 5. Cryo-EM reconstructions of the P7-1 contracted sheath determined at pH 4.5.** Atomic models of the N-terminal portions of LTF (orange) and STF (cyan) are overlaid onto the experimental density calculated at 4.2 Å resolution.

**Supplementary Table S1: Sequence of the predicted structural proteins of P7-1**

| name | size (aa) | sequence |
|---|---|---|
| **gp67** | 480 | MAETEKTAPGIPRLRLGEIGSTGLKQVNGTILEERRPELRFPRACRTFQMMAEDPTIKSAL<br>DLFEMMMSRVDWEVDLGVDPDEAMKARGKFLKECMHDMEHSWYSFIKEVTSFYTYG<br>FSVHEMVFKRREGYPVSKYNDNKFGIKKLPIRSQSTITKWLYSEDGRNFLGCEQSLANVV<br>NGDRYVNLGNKDGTVEIPAKKMLLFRVDAKRDSPEGNSPLRACYNAWRYRVEIEEQES<br>VGVTRDMNGMPTLYLPPRYMSEDATEAERAVYEYYKRVIRNIQMNEQSGLILPQAFDP<br>ESRQPLFKFELTSSQGSKMYDTDAIIRRWDNKILQALFADMLKMGQDQVGSYSLAGAK<br>TNIMAMAIESRLREIKDVLDNKLIPTLFALNGDYSPDLPKLQYGELDEIDLEEFSKGIQRIGS<br>VGGLERDREVYNKIRKALKIKPRPDDEPVDVDNIMGGQSQAGKAGVGNGASTSAAGR<br>DNAAANNA |
| **gp70** | 137 | MAQNIIAKDHQRLSNWLKEEQMGHRGLFYTRETLPVADIDVKTTGSVLDSTGKLVTKAT<br>IADATYILITDLHDYANAQMSHAVVLARGFAKIGSKAVIFGADVDDADKATVFEAFKAKN<br>IFAVDQIEGAFENVTFA |
| **gp71** | 345 | MANTRSYLNDGQFYIADQTENLLIIPNTWTLVENMGVFTSEGVTQNTVQFEEIETRYGL<br>VKDAIRGTRHQVASDQRRQLRAFAIPHFNQDDYITPEDIQGKRAFGADREETLNEVRAR<br>KLETIRRNWANTAEVASVSAIVTGKSYAPAGTIEYDWYDLMGKTRKVVGFDLTNPTADV<br>MGKTEEIFVHMQDNSQDGLIRGDFVALCSPEFFTALINHPSIKEFYKAYQASPQYWRERL<br>TARGLDLRFREFYFGNIHFIEYRGVDPYGNRLIPAGDAYFIPTDSGDLFARYFGPGSTFDDL<br>GTLGKELYATERMAEDRRSILIETESNFIHVLRRPQMIVRGTVNA |
| **gp72** | 159 | MAYTGDPANNPIDRLREIVGDVWEPPMLSDETYQWVLDKNEGNERRAALELMRMML<br>FRLTRGMRERTGDIEVYGAEYFNNYLKALQLILKDPNIAISLAVPYAGGISKSDMLANDLD<br>PDNVTREFYIGFARREKLYNQCNPGPQDLSMGCDYLGKLQF |
| **gp74** | 127 | MLYPTFSMTNFVKLDLIRRGQPGDDGFGRPTPPVETVVTITANVQPIEKSTDTRILPEAD<br>RSKACFKVYSRGEEIRQLKEGPGGWSADRFMWEGELYEVMKVINYSMGILNHYKAICM<br>RVERNSTA |
| **gp75** | 128 | MIHTELENSLYLMVKELFPDWRVIQAYTNNQEPQTPYLAIDIKRLDELGRENVSNMSDPI<br>SPSHGTIQVQQDFAAKVVFELIGKYGETASVSDMAMAITRAMRTPAGHAAQRKFNLSL<br>FKLPSTRRVPMLRETDMYMFYQVTCEFGFSVIETTTQEFAAGADIHGVYYDAGRPGHIIE<br>SHIDINFEH |
| **gp76** | 429 | MTVLTDVIDIQISRETAAVAQTNFNVPLFIASHTNFSERARVYNSLKGVAEDFGESDPTYL<br>AAVRYFGQALKPRSLVIGRRQVPSATVSVSVVQEGQSYVLTVNGLPVSYVSQQDDTATLI<br>ATGLKAAYDVTPVAGVTVTDNEDGTLTVASNEDWSLKVSSNLTMAAAPSTEGWPAAIT<br>AVQGENDEWYALSIDSHADDDIMAVATHIEGTKKVFIGATAQANTKTSAENDIASRLVA<br>AGFQRTALIYHPNADAQFPECAWVGYQLQEQPGSNTWTHKALAAVDAYRLTPTESTNL<br>KNKNVTTFERVGGVNRTFGGAMAGGEWIDVMIFVDWLEARMTERLWFRMANSKKIP<br>YDAVGATILESEIRAQLNEGIRVGGLAEAPAPKVFVPDVLSMSPNMRAQRIFEGIEFEARL<br>AGAIHFVHIRGTVTV |
| **gp77** | 175 | MAVQRLATFSPADVTIVITHPATGESMVLGGFSEDSIVNIERNADTYVMYTGADNTSTR<br>VYNASKSATLTVSLQQTSPSNDFLTALYNYDDARRSSEGLFTIHVKDNSGRSDYFSDDAY<br>VGVVPGSNFSNSMQTRDWVIHAHNLQTLIGGNAKLSPGDADTLRNLGVTLDQRWL |

| | | |
|---|---|---|
| **gp78** | 167 | MITTYSPRDVVVTLAGIHSVTGYAEGEFIRIVKDIKPFIKHGSMDGEIARVYNKDQGWRV<br>ELTIMQSSPTNDILSMLYNVDIATRMGKFPLMIKDTKGSTSFLALTAWVEDLPRVSFSGQ<br>LETRTWILGCSEVAMNIGGNVDQSLVEQAILLGSSLLPALTQFGGF |

| name | size (aa) | sequence |
|---|---|---|
| **gp79** | 160 | MANTVLTYSPSDVKIVLCGYALTGVVSFEMSWLSRPYTMVRGIRGHHTRVFNRDLSA<br>QIRIEVLQTSVSNDAFFSLVEQDRRTQSARITLSVKDTHGSTMMSTDNAYVNGYPSITF<br>TDGIENRVWTIDVLDWTDGTVGGNQQVGFDVFGTVQGALSYLR |
| **gp82** | 789 | MITERIAQLTGELKFTVDSRPLTAFDKKLAGVEARLREFSKLTNKRFGVKLTLDTKTLREE<br>LAKAATQRVVLKNIAVDAAAVRLISEKLQERLNATPIRLKSVRLDLSGIRDQKNFVKTAL<br>GQTKVDLPVELGLAQASRTLYEWKKRTESRFKIHLNADISRSKLLQNARNTLRDVQGRL<br>NGLAVATPQIRLSVDRAHLRREIQDVLEQIRREVRIRIDLESSIRGGGAGGSRGTAGHIR<br>QGMGMGIGSELAGWGRGFIPGLGGAFAIMQLNRANQELQGQRLAMQAVGGGVQ<br>GGQELQATLRDISQRLGLDDRAIGSSYVKMMAAGQASNFDKTQVDGIFQSMAEYGR<br>VMGLDGEAMKGSFRAVEQMMGKGQIMSEELKGQLAERFPAAVALMAKSQDMTIA<br>ELMKTMEKGELKSDALIPFARTLAEEARKGGALDAAMQGTAAQQGRFQFGWNRTIE<br>AFAAGGFDRGMSDFFKIAAQGMKEALPLVTALGGGFEALMRPVNALVGIGGELGSQ<br>WENIAKQFNMTGTGLTLTTAQTLALLTPMGRLISAISWGALALEDFIVFLEGGDSVFG<br>DFLNNNVQAAETFEKLASESSELKNNLDGIFSVVPGLAEALKGLEFNEMLVSTMRELA<br>AIMEFFNSVVERFAIAGKYRDAKIAEAGGDRSTIMSNIDTMYAMFNPEDAKDKATFIG<br>DRLVAQNIDAIQTETHTRATQSLTPDQFGYMMQRGQVRNEMEGALKKSAIDISFNLN<br>VSGIDAQGNVMTTEAQERVREIVSNVLEEEISRASASYKESQ |
| **gp83** | 254 | MTIAIKRENGDLIWFDAVTEFGRQYRGSVSSNPIETGGKITDHITTENPVFTLTAVVSD<br>ADFNLNRPVINDNEAQTYKINNKEFVNTQPVTIPAVISTSRLDPSRIFPEVITQFIPPEIPS<br>ATVLPQKGDKVAYDIERQLIDMQRNAEVFSLLDFRDGIIWDQIERCIFTDLSFTENAET<br>GSSLQPRMTIEAVTFTDTRYVEVRVNKGRKTAKKEKRDTKEGDTGASNATSQDSPTFK<br>RSQMKEAQIRTQVAR |
| **gp84** | 119 | MSTTYIDTLPLYQDRKYRYAVAIEGISRVLQFYWNSRSRQWHMDIFDEELNPILTGLAV<br>VPQYPIMADYAMQHIGFNGYFLLMPVNLEQVQYKHDASDIVPQFFELLYVNVDLEDD<br>VE |
| **gp85** | 306 | MKQYDRVYKLTLGNTESGQGVEITNLHPDGSLNREGLQFRFDISKSSDNKKSGNSATV<br>EIYNLSIATLNILETEYLTCRLEVGYKEMGTSVVLDGNVVETSTRKSGNDYVTQLILGEGY<br>TALTETKLKGTVSPGKTVKDVIEEIRLQMPGVDRGAYTGLNCNNPIMYGYQLRGLAKD<br>ALNSVCEANNIEWNISGNVLNVTDVNGPTTKSVQMAPLVNRETGLIDIPFYASASGTA<br>QKKDQRRRRGVQFKCLLNPELTPGVLVRVESDRLSGTFRINNVRISGGYRDNEWYTEC<br>WCSDLNQEDIDQ |
| **gp86** | 247 | MRRTGLQELLNLHSSTEGSKQYTAIPCVVLRVLDDFRRLSVDVRPVVNDLYKDGTSEE<br>QPEILSVPVIMPGTANTLISFPLNVGDTVLCVFSQRTMDVFKGSATGQPHTPNDLRKF<br>NMADAIAIPGLFTFPRSMNDPARHSWPHDTKDLTIAHNLMTGQEAEVRIKANGDILI<br>NSRKTISINAQNVNVKCASLTVDAAHTTWNGNIAHTGNYVQTGGTSTFNGIPFHTHK<br>HGGVMPGGGVTAVPQA |

| | | |
|---|---|---|
| **gp87** | 124 | MDLFVNPDTHDLVFINGEAPVTQRMVDIVAQRLKIKLYTFLGEWFLDDRIGIPYFERILGKSRSLPAVDAIFQSEIMRDPGVLEITSWQSGIDPHTREYSMEFTVRTTDNTESLPITFRMIGV |

| name | size (aa) | sequence |
|---|---|---|
| **gp87** | 124 | MDLFVNPDTHDLVFINGEAPVTQRMVDIVAQRLKIKLYTFLGEWFLDDRIGIPYFERILGKSRSLPAVDAIFQSEIMRDPGVLEITSWQSGIDPHTREYSMEFTVRTTDNTESLPITFRMIGV |
| **gp88** | 488 | MAGITAEGLTIKDLDQILTDYRNVASQVFADLVSTGDEVDTSGNSALGRLIGVVAPSDEAIWEVIQMVYNSFNPAAATGVALDNLVSFSAISRHAARPTRAQVVLEGNINTVINSPPAKMISSSTGRVFHLLQGVILTPKACSGVGIFPQTVGNDLTYELKMYVDDINTTSIKYTSPATGTVTSESILAGLAADVATNHGNTLTSYEQNGILFIVPIDPFNTKTFEEASNLSIQKVRKLGIAVDDVIGPVPQQALAIDTISIPIAGWDSVINPVPAITGRLRETDDELRERFRNSKFVQATNILESLIDGLMNVEGVEDVRIIENDTDNPDPVHGVPGHSFLPIVLGGIPTEVAQSIWLNKPFGIGSVGDTEVQVVDSRGYTHRVNYQRPVEVPIEIKISVTNTGSMPENIEDILRPRIVAYGTENYKIGDDVIYSRFYAPIMEIPGFQVNSLTIAKKGQTQGMANIEIGFKEVATFAAADITVTTV |
| **gp89** | 244 | MAVNQFDREDYLEVARERVTEQFKEKPIFDRFLQVLLSGKFDIQNALEDLQTLRSLDTATGKQLDIIGDIVGRPRGLVYQDIFNYFGFAGTERAGSFGSLSDPTVGAPWYSVGAPTGNAREPSDEEYRMILKAKIIKNRTNSTPEQVIEAYKFVFGVPEVFLEEYAPAAVRIGIGKILTNVERSLLFDLGGAGALLPKTIGVNYTYTEFQAGRVFATEGFPGGQGVGDLNDPTVGGILTNLVT |
| **gp90** | 672 | MADYSQLPIENIWSTGGDMVAPTPAQQQGGWGIQSVPRQWWNWKWNLHDTNLAYLLQKGIPEWTSTQEYIANKSFCTRGGFVYKAVRTHTGSDPAQVSANWVRAFADYTTSSSALGSLTPREGGIPFFISATGASVFDSTAYGRGMLNVANASAARSYISAQESSVVLSNLSTVTRAANTVPYFNTDTSMATFNITAFGRGLVNAANDENARNFLGLANSAIITADPANRAHTLVYRDAAGNFNAGVITATLSGNATTANKLRTPVTINGVAFDGSQNIVLPGLDTSYAGTVARLHINGANLSSADKTTQLALRNKSDNDWISLAVVDDNILQFVFRSATNPVVQIGNEVILHTGNQFSLGPTLTDARSRLGLDRLTQGSSDTQVFPSTLNNGPYLTVQPTAIGGFNGSTNGWLFRFDANGNMTHGTVPAARITGLSNSAQIPATTTAQANSLVQRDADGGFSAIAINAYGTIVGYGSNIFSRASGTGNAHIGFQRANGTELGLIWGAQSNNSMNFRVAGGATAVSITGLDMTVTGRVNATTLNASGNVNATGNVNAGSATLNTAGNITGAAYGAYGSLTNWVDSVYAKKGEIPNDIARAGAAWDAVGQYILAGDQSGGSGGPGTTRAGSQLKPYSTISYTAGALPAAGTYRCMGAFAGGGNQITLWQRIS |
| **gp92** | 500 | MPNIMKPTGINAIWSENGQKVDPGAVKVGLGWVTELPPYQTANFIEYKQDLFNAHVNQHGIPEWDSVTEYQGNLSYTQGANGIIYKCLRTHSDKIPTDPLNITAGYWRVAFEDAGEAAKVQANLDRHVTNYNTLSGIGNVVIARQNLSVYSKAEGDARYAMKHGNGSNVFSVATATQPTHAIPLSQLSTLVPPATETVAGVMAVATTIETEAGANDTKAVSPLKAAQVYLKKKDNLSGLSNVTAARANLGLSDTATMPSSTFLKAGSNLADVPNKALARSNLGITSSATQPETYFLRSAQNLADVPNKAQARVNLGLTGMATTDPAAVMMKADNLAGLANTATARSNLGLGTASTRNTGDFLSSGDNLSDLTNVQAARNNLGLKGAATLDVWGLPANTTAMDFQSNQSDISRGWARLPNGLLLQWGTGPGLSDDTRTNIQLPVPARILNVQVTVMGTFNNSIGPGAFMTDMWSNTGFRVSCNWGNWSYPFNWFAITSTL |

**Supplementary Table 2: Identification of P7-1 Major Structural Proteins by LC-MS/MS**

| Protein | Mascot Score[a] | Mass | Molecular Weight (kDa) | Annotation | Predicted size (aa) |
|---|---|---|---|---|---|
| gp71 | 9035 | 39407 | 39.4 | putative major capsid protein | 344 |
| gp90 | 5993 | 70018 | 70.0 | putative tail fiber protein | 671 |
| gp67 | 4786 | 54486 | 54.5 | unknown | 479 |
| gp82 | 3824 | 85861 | 85.9 | Putative tape-measure protein | 788 |
| gp88 | 2813 | 52411 | 52.4 | putative baseplate protein | 487 |
| gp92 | 2789 | 53142 | 53.1 | putative tail fiber | 499 |
| gp70 | 2453 | 14841 | 14.8 | unknown | 136 |
| gp76 | 2087 | 46353 | 46.4 | unknown | 428 |
| gp77 | 1927 | 18951 | 19.0 | unknown | 174 |
| gp83 | 975 | 28582 | 28.6 | unknown | 253 |
| gp72 | 777 | 18227 | 18.2 | unknown | 158 |
| gp85 | 628 | 34168 | 34.2 | unknown | 305 |
| gp75 | 526 | 21326 | 21.3 | unknown | 187 |
| gp69 | 510 | 33046 | 33.0 | unknown | 305 |
| gp89 | 463 | 26658 | 26.7 | unknown | 243 |
| gp87 | 410 | 14173 | 14.2 | unknown | 123 |
| gp86 | 378 | 26836 | 26.8 | putative baseplate protein | 246 |
| gp79.1 | 195 | 17745 | 17.7 | unknown | 159 |
| gp78 | 193 | 18294 | 18.3 | unknown | 166 |
| gp84 | 167 | 13983 | 14.0 | unknown | 118 |
| gp68 | 146 | 17265 | 17.3 | putative methyl transferase | 156 |
| gp74 | 145 | 14456 | 14.5 | unknown | 126 |

[a] Statistical measurement of how well the observed MS data matches the identified protein sequence; scores greater than 50 represent correct identifications and can reflect relative protein abundance within a sample.

**Supplementary Table 3: List of *Pakpunavirus* genomes used in this study**

| Phage name | Accession Number | notes |
|---|---|---|
| 20Sep416 | NC_073612 | |
| 20Sep418 | OQ319937 | |
| 908-1 | OQ319933 | |
| Ab15 | LN610587.1 | Annotated for this study |
| Baskent_P3_3B | PP766723 | |
| bmx-p3 | OQ319933 | |
| C11 | NC_028652.1 | |
| EM | NC_073601 | |
| GEC_MRC | PP836135 | |
| GEC_PNG3 | PP836136 | |
| Henu5 | NC_073608.1 | |
| HJ01 | NC_073615 | |
| ITTPL | NC_073605.1 | |
| JG004 | NC_019450.1 | |
| K5 | NC_030910.1 | |
| K8 | NC_028817.1 | |
| Kat | OQ992554.1 | |
| LPS-5 | PP203294 | |
| PA10_SL - PA10 | NC_041903.1 | |
| PA45_GUMS | MN563785 | |
| PaGz-1 | MH791399 | |
| PAK_P1 | NC_015294.2 | |
| PAK_P2 | NC_022967.1 | |
| PAK_P4 | NC_022986.1 | |
| PaoP5 | NC_029083.1 | |
| PaP1 | NC_019913.1 | |
| PaSz-1_45_92k | MN871481 | |
| PaSzW-1 | MH791416 | |
| Pa-U | MN871470 | Annotated for this study |
| PaYy-2 | MH725810 | |
| PaZq-1 | MH791408 | |
| phiMK | NC_031110.1 | |
| phipa10 | OK539826 | |

| | | |
|---|---|---|
| PhL_UNISO_PA-DSM_ph0034 | MW526259 | |
| PIAS | SRR14274931_1 | Assembled and annotated for this study |
| PJNP029 | OR941787 | |
| pPA-3099-2aT.2 | NC_073606 | |
| PseuPha1 | OP756059 | |
| SCUT-S2 | MK340761 | |
| SPA01 | NC_073602 | |
| SPA05 | NC_073604 | |
| SRT6 | MH370478.1 | |
| vB_PaeM_B31 | NC_073614 | |
| vB_PaeM_B55 | NC_073618 | |
| vB_PaeM_C2-10_Ab02 | NC_042113 | |
| vB_PaeM_C2-10_Ab1 | NC_019918.1 | |
| vB_PaeM_C2-10_Ab10 | LN610586 | Annotated for this study |
| vB_PaeM_MAG1 | NC_031073.1 | |
| vB_Paer_Ps12 | NC_073621 | |
| vB_Paer_Ps25 | OM870968 | |
| vB_Paer_PsCh | NC_073623 | |
| vB_Paer_PsIn | NC_073622 | |
| vB_PaeS_B8 | NC_073616 | |
| vB_PaM_EPA1 | NC_073609.1 | |
| vB_VIPPAEUMC01 | OQ721915 | |
| vB-PaeM-YQ78 | OP627531 | |
| vFB297 | OQ921398 | |
| YS35 | NC_073617.1 | |
| Zigelbrucke | NC_041904.1 | |

**Supplementary Table 4. Variability in the sequence of LTF (gp90) homologs within and across the four groups.** The table shows the protein sequence identity for LTF among phylogenetically related *Pakpunaviruses*.

| | 7-1-like | PAK_P1-like | EM-like | C11-like |
|---|---|---|---|---|
| **7-1-like (n=30)** | 95-4-100% | 86.2-89.6% | 72.6-73.2% | 64.7-69.1% |
| **PAK_P1-like (n=3)** | | 93.3-98.8% | 73.5-73.9% | 65.6-69.1% |
| **EM-like (n=1)** | | | 100% | 67.1-69.1% |
| **C11-like (n=27)** | | | | 91-100% |

**Supplementary Table 5: List of phages used in the analysis of LTF sequence conservation presented in Figure 7**

| 7-1-like | | | |
|---|---|---|---|
| vB-PaeM-YQ78 | OP627531 | PAK_P2 | NC_022967.1 |
| HJ01 | NC_073615 | phipa10 | OK539826 |
| vB_VIPPAEUMC01 | OQ721915 | vB_PaeM_C2-10_Ab08 | LN610575.1 |
| PA45_GUMS | MN563785 | Ab15 | LN610587.1 |
| pPA-3099-2aT.2 | NC_073606 | vB_PaeM_C2-10_Ab1 | NC_019918.1 |
| vB_PaeM_B31 | NC_073614 | vB_PaeM_C2-10_Ab10 | LN610586 |
| vB_PaeM_C2-10_Ab02 | NC_042113 | vB_Paer_PsCh | NC_073623 |
| PIAS | PRJNA722489 | PA10 | NC_041903.1 |
| vB_PaeS_B8 | NC_073616 | PaP1 | NC_019913.1 |
| YS35 | NC_073617.1 | K5 | NC_030910.1 |
| GEC_PNG3 | PP836136 | K8 | NC_028817.1 |
| GEC_MRC | PP836135 | PAK_P4 | NC_022986.1 |
| vB_Paer_Ps12 | NC_073621 | PJNP029 | OR941787 |
| vB_Paer_Ps25 | OM870968 | vB_PaeM_B55 | NC_073618 |
| vB_Paer_PsIn | NC_073622 | | |

| PAK_P1-like | |
|---|---|
| vB_PaeM_MAG1 | NC_031073.1 |
| PAK_P1 | NC_015294.2 |
| ITTPL | NC_073605.1 |

| EM-like | |
|---|---|
| EM | NC_073601 |

| C11-like | | | |
|---|---|---|---|
| Pa-U | MN871470 | PaSzW-1 | MH791416 |
| SCUT-S2 | MK340761 | PaZq-1 | MH791408 |
| vB_PaM_EPA1 | NC_073609.1 | PseuPha1 | OP756059 |
| phiMK | NC_031110.1 | Kat | OQ992554.1 |
| PhL_UNISO_PA-DSM_ph0034 | MW526259 | SPA05 | NC_073604 |
| Zigelbrucke | NC_041904.1 | PaGz-1 | MH791399 |
| 20Sep416 | NC_073612 | SPA01 | NC_073602 |
| PaoP5 | NC_029083.1 | C11 | NC_028652.1 |
| 20Sep418 | OQ319937 | Henu5 | NC_073608.1 |
| 908-1 | OQ319933 | SRT6 | MH370478.1 |
| Baskent_P3_3B | PP766723 | PaYy-2 | MH725810 |
| bmx-p3 | OQ319933 | vFB297 | OQ921398 |
| LPS-5 | PP203294 | JG004 | NC_019450.1 |
| PaSz-1_45_92k | MN871481 | | |
